# How to make a rodent giant: Genomic basis and tradeoffs of gigantism in the capybara, the world’s largest rodent

**DOI:** 10.1101/424606

**Authors:** Santiago Herrera-Álvarez, Elinor Karlsson, Oliver A. Ryder, Kerstin Lindblad-Toh, Andrew J. Crawford

**Author notes:** **Corresponding author**: Address: Universidad de los Andes, Calle 19 No. 1-60, Bogotá, código postal 111711, Colombia, South America.

## Abstract

Gigantism is the result of one lineage within a clade evolving extremely large body size relative to its small-bodied ancestors, a phenomenon observed numerous times in animals. Theory predicts that the evolution of giants should be constrained by two tradeoffs. First, because body size is negatively correlated with population size, purifying selection is expected to be less efficient in species of large body size, leading to a genome-wide elevation of the ratio of non-synonymous to synonymous substitution rates (*d*_N_/d_S_) or mutation load. Second, gigantism is achieved through higher number of cells and higher rates of cell proliferation, thus increasing the likelihood of cancer. However, the incidence of cancer in gigantic animals is lower than the theoretical expectation, a phenomenon referred to as Peto’s Paradox. To explore the genetic basis of gigantism in rodents and uncover genomic signatures of gigantism-related tradeoffs, we sequenced the genome of the capybara, the world’s largest living rodent. We found that d_N_/d_S_ is elevated genome wide in the capybara, relative to other rodents, implying a higher mutation load. Conversely, a genome-wide scan for adaptive protein evolution in the capybara highlighted several genes involved in growth regulation by the insulin/insulin-like growth factor signaling (IIS) pathway. Capybara-specific gene-family expansions included a putative novel anticancer adaptation that involves T cell-mediated tumor suppression, offering a potential resolution to Peto’s Paradox in this lineage. Gene interaction network analyses also revealed that size regulators function simultaneously as growth factors and oncogenes, creating an evolutionary conflict. Based on our findings, we hypothesize that gigantism in the capybara likely involved three evolutionary steps: 1) Increase in body size by cell proliferation through the ISS pathway, 2) coupled evolution of growth-regulatory and cancer-suppression mechanisms, possibly driven by intragenomic conflict, and 3) establishment of the T cell-mediated tumor suppression pathway as an anticancer adaptation. Interestingly, increased mutation load appears to be an inevitable outcome of an increase in body size.

**Author Summary:** The existence of gigantic animals presents an evolutionary puzzle. Larger animals have more cells and undergo exponentially more cell divisions, thus, they should have enormous rates of cancer. Moreover, large animals also have smaller populations making them vulnerable to extinction. So, how do gigantic animals such as elephants and blue whales protect themselves from cancer, and what are the consequences of evolving a large size on the ‘genetic health’ of a species? To address these questions we sequenced the genome of the capybara, the world’s largest rodent, and performed comparative genomic analyses to identify the genes and pathways involved in growth regulation and cancer suppression. We found that the insulin-signaling pathway was involved in the evolution of gigantism in the capybara. We also found a putative novel anticancer mechanism mediated by the detection of tumors by T-cells, offering a potential solution to how capybaras mitigated the tradeoff imposed by cancer. Furthermore, we show that capybara genome harbors a higher proportion of slightly deleterious mutations relative to all other rodent genomes. Overall, this study provides insights at the genomic level into the evolution of a complex and extreme phenotype, and offers a detailed picture of how the evolution of a giant body size in the capybara has shaped its genome.

## Introduction

Body size is arguably the most apparent characteristic when studying multicellular life, and understanding how and why body size evolves has been a major question in biology [1]. Body size is a canonical complex trait, and, consequently, ontogenetic changes underlying body size evolution affect other life-history traits, such as fecundity and longevity [2,3]. Yet, how size is developmentally controlled to generate species-specific differences remains elusive. Although increase in body size has several selective advantages [4], maximum body size in tetrapods appears to be determined mostly by intrinsic biological constraints [5–7]. Two major tradeoffs in the evolution of large bodies are the reduction in population size and the increased risk of developing growth-associated diseases, such as cancer [8,9].

Larger species have lower population densities [8], slower metabolic rates, longer generation times, and a lower reproductive output [3]. Together these traits lead to an overall reduction in the genetic effective population size (*N_e_*) relative to small-bodied species [10]. In a finite population, the interplay between selection and genetic drift determines the fate of newly arisen mutations, and the strength of drift relative to selection will largely depend on *N_e_* [11]. In smaller populations, slightly deleterious mutations are expected to accumulate at a higher rate than in larger populations, due to stronger genetic drift relative to purifying selection, thus increasing the mutation load in the population, which may increase the risk of extinction [12]. Furthermore, animals have positive allometric scaling of number of cells in the body with overall body size [13]. Thus, if every cell has an equal probability, however low, of becoming cancerous and related species have equivalent cancer suppression mechanisms, the risk of developing cancer should increase as a function of number of cells. Therefore, large species should face a higher lifetime risk of cancer simply because large bodies contain more cells [9,14]. Moreover, giant body sizes are achieved mainly by an accelerated post-natal growth rate [15], which represents an increment in the rate of cell proliferation [16]. For a given mutation rate, an elevated rate of cell proliferation accelerates the rate of accumulation of mutations and, thus, increases the chances for a cell population of becoming cancerous [17].

The continued existence of enormous animals, such as whales and elephants, implies, however, that the evolution of gigantism must be coupled with the evolution of cancer suppression mechanisms. In fact, the lack of correlation between size and cancer incidence, known as Peto’s Paradox, suggests that large-sized species have mitigated the inherent increase in cancer risk associated with the evolution of large bodies [9,14]. For instance, genomes of the Asian and African elephants (*Elephas maximus; Loxodonta africana*) and the bowhead whale (*Balaena mysticetus*), three of the largest living mammals, revealed lineage-specific expansion of genes associated with tumor suppression via enhanced DNA-damage response [18,19] and cell cycle control [20], respectively. Thus, cancer selection, i.e., selection to prevent or postpone deaths due to cancer at least until after attainment of reproductive maturity, is especially important as animals evolve larger bodies and might account for the relative lower incidence of cancer in large animals and the evolution of anticancer mechanisms [21].

Among mammals, rodents are the most diverse group in terms of species richness and morphological disparity, particularly in body size. South American rodents of the infraorder Caviomorpha represent one of the most spectacular evolutionary radiations among living New World mammals [22], and fossil data suggest the independent evolution of gigantism in at least three lineages [23]. Caviomorphs have the broadest range in body size within rodents, and make Rodentia one of the mammalian orders with the largest range in body size [24].

Most notable among the caviomorphs is the largest living rodent, the capybara, *Hydrochoerus hydrochaeris* ([Linnaeus, 1776]; Fig. 1C), family Caviidae. Capybaras have an average adult body weight of 55 kg [25], being 60 times more massive than their closest living relative, the rock cavies (*Kerodon* sp.), and ~2,000 times more massive than the common mouse (*Mus musculus*). Relative to all rodents, capybaras are three orders of magnitude above the mean body size for the order [25]. Previous studies on physiology and molecular evolution identified that caviomorph insulin, a hormone involved in growth and metabolism, has a higher mitogenic effect relative to other mammalian growth factors [26,27], suggesting that lineage-specific changes in insulin within caviomorphs coincide with important changes in life-history traits. However, the genomic basis of the capybara’s unique size remains unexplored. Additionally, there is evidence that telomerase activity has co-evolved with body mass in rodents, with the largest species (i.e., capybara and beaver) displaying low telomerase activity in somatic tissues, suggesting a potential role for replicative senescence as a cancer suppression mechanism in large-bodied rodents [28]. However, it is yet not clear whether capybara has a unique solution to its increased risk of cancer relative to smaller rodents, or how some lineages have coupled growth-regulatory pathways and cancer suppression mechanisms to permit evolution of giant bodies.

**Fig 1.**
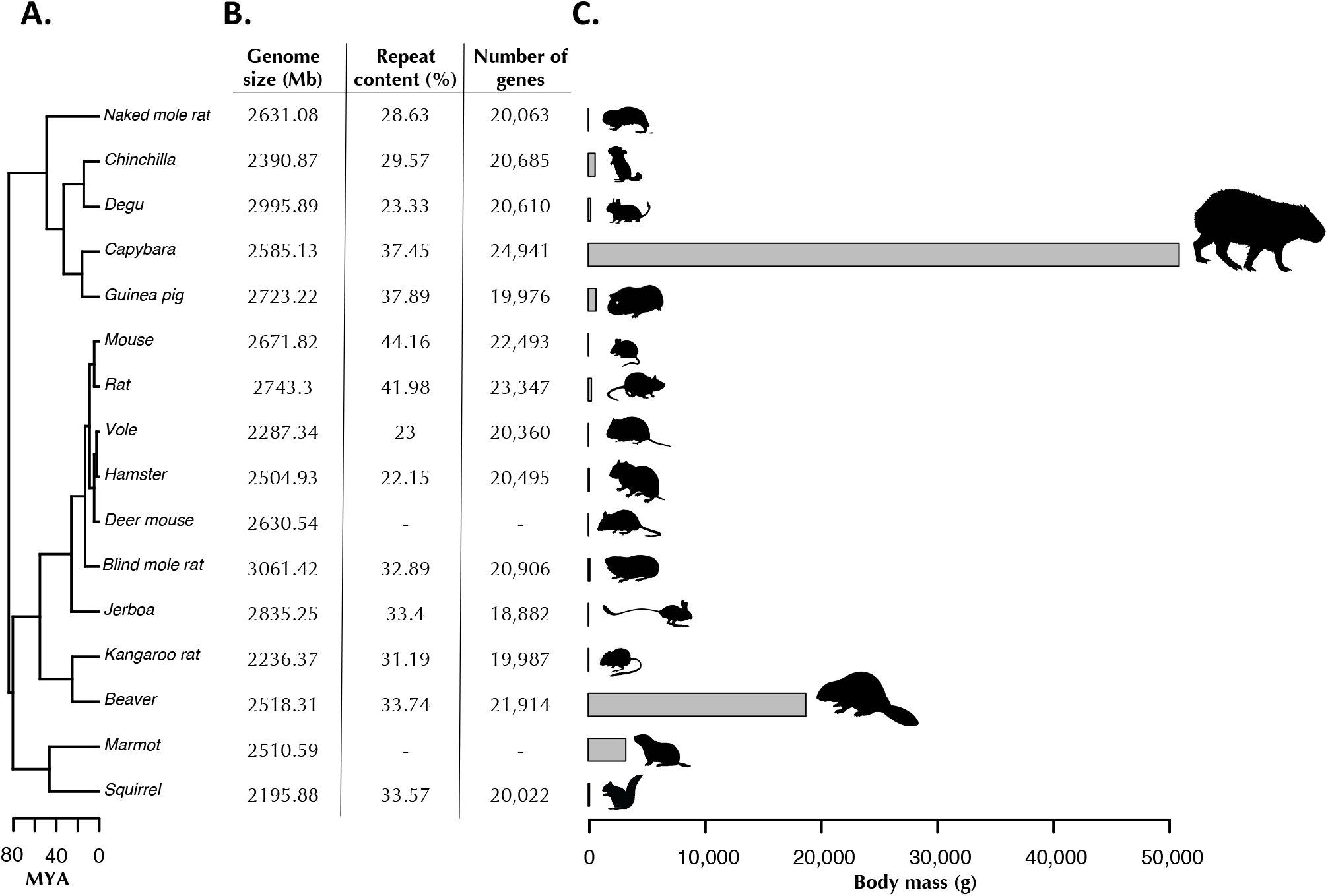
Divergence times, genome statistics and body mass of representative rodents used for comparative genomics analyses. **(A)** Divergence times of rodent species using the topology obtained from the phylogenomic analysis (see Materials and methods). MYA: Million years ago **(B)** Basic genome statistics of rodents used for comparative analyses. Neither Repeat Content nor Gene Content was available for Deer mouse and Marmot **(C)** Body mass ranges from 20.5 g, being the mouse the smallest surveyed rodent, to 55,000 g in the capybara, the largest living rodent.

## Results

### Genome landscape and annotation

Capybaras have 32 autosomes (2*n* = 66), consisting of 10 submetacentric chromosome pairs, 10 metacentric chromosome pairs, 2 acrocentric chromosome pairs and 10 telocentric chromosome pairs [29]. We generated a whole-genome sequence and *de novo* assembly of a female capybara, using a combination of a DISCOVAR *de novo* assembly and Chicago libraries (See Materials and methods). The total length of the assembly was ~2.7 Gb, slightly shorter than the genome size estimated by flow cytometry of 3.14 Gb (Table 1), including 73,920 contigs with an N50 of 12.2 Mb and the longest contig at 75.7 Mb. The GC content was estimated at 39.79%.

**Table 1.** Assembly statistics of the capybara genome.

|  | **Starting Assembly**<br>**(Shotgun + DISCOVAR *de novo*)** | **Final Assembly**<br>**(Chicago + HiRise)** |
| --- | --- | --- |
| Total length (Mb) | 2,735 | 2,738 |
| Contig N50 (Kb) | 148 | 161 |
| Number of scaffolds | 656,018 | 73,920 |
| Scaffold N50 (Mb) | 0.202 | 12.2 |

The assembled capybara genome has very high coverage of coding regions: we recovered 89% of the 3,023 vertebrate benchmarking universal single-copy orthologs (BUSCO pipeline, [30]), and 94.4% of the ultra-conserved core eukaryotic genes (CEGMA pipeline, [31]), which is comparable to the guinea pig genome despite the latter having two-fold larger scaffold N50 (Fig. S1A). We used guinea pig, rat, and mouse Ensembl protein sets to guide the homology-based annotation with MAKER v2.31.9 [32], resulting in a capybara genome containing 24,941 protein-coding genes with 92.5% of the annotations with an AED bellow 0.5 (See Materials and methods). RepeatMasker annotation and classification of repetitive elements estimated that 37.4% of the capybara genome corresponded to repetitive content, spanning 1,025 Mb of the genome (Fig S1B). Most of the repeat content (72.5%) corresponded to transposable elements, with LINE-1 the most abundant (20.46%), similar to the mouse genome [33]. Further comparison across the rodent phylogeny showed that both genome size and repeat content of the capybara are comparable to that of other rodents (Fig. 1B).

### Body mass evolution

To determine whether the capybara could be considered a giant based on a lineage-specific accelerative rate of body mass evolution, we used phylogenetic comparative methods. We found that the rate of body mass evolution among caviomorph rodents has not been constant throughout the their phylogenetic history, as shown by the better fit of a multi-rate model compared to the single-rate-Brownian Motion model (ΔDIC = 78.6; Table S1) with an estimated value of α = 1.35 (95% confidence interval [CI]: 1.0 to 1.7). The ancestral body size of the most recent common ancestor (MRCA) of Caviomorpha was estimated to be 971 g (95% CI, 221 g to 5,135 g) and the size of the MRCA of the capybara and rock cavy (*Kerodon rupestris*) was estimated to be 1132 g (95% CI, 437 g to 23,225 g), suggesting that the rate of body size evolution has undergone one or more sudden changes.

Using the AUTEUR package [34], we detected three shifts in the rate of body-mass evolution along the caviomorph tree (Fig. S5): one decrease in evolutionary rate at the root of the Octodontoidea clade (shift probability: 0.11), and two incidences of rate acceleration. The first rate increase was localized on the branch leading to the coypu (*Myocastor coypus*; shift probability: 0.16), the largest species within Octodontoidea, and a second increase, with the highest posterior support, in the MRCA of the capybara, rock cavy and Patagonian mara (*Dolichotis patagonum*; shift probability: 0.44). Capybaras and Patagonian maras are the largest living species of caviomorph rodents, yet the capybara is approximately 3 times heavier (Fig S6). These results suggest that the capybara evolved from a moderately small ancestor, comparable to the size of a guinea pig, indicating that capybara’s characteristic large body was achieved by a spurt in the rate of body mass evolution (see also [35]).

The accelerated rate of body mass evolution observed in capybaras relative to other rodents parallels the rate acceleration observed in the world’s largest animals, e.g., elephants, the largest living terrestrial mammals relative to smaller afrotherian ancestors [36], baleen whales including the blue whale, the largest animal to have ever lived, relative to ancestral mysticetes [37], and sauropodomorphs, the largest terrestrial animals in Earth history, relative to other dinosaurs [38]. Thus, the tempo and mode of body size evolution demonstrates that the capybara also provides an example of gigantism among rodents.

Having characterized the genome and the mode of body mass evolution, we undertook detailed analyses of key genes and gene families to explore the genomic basis of capybara’s large size, identify possible molecular adaptations related to cancer suppression, and detect the genomic signatures associated with a reduced *N_e._*

### Gene-specific rates of evolution in the capybara genome

Because the guinea pig (*Cavia porcellus*) is the closest rodent with a sequenced genome to the capybara (Fig. 1A) and is much smaller (750 g versus 55,000 g), pairwise comparisons between the capybara and guinea pig could provide insights into the genes underlying the rapid burst in body size evolution in the capybara (see *Body size evolution*). We calculated the pairwise d_N_/d_S_ ratios for 11,255 single-copy gene orthologs (SCOs) shared by capybara and guinea pig. Sequence conservation in protein-coding genes between guinea pig and capybara was moderately high (mean = 92.6% with standard deviation [SD] = 4.8%) after an estimated divergence time of ~20 million years (Fig. S3). The 20 capybara-guinea pig SCOs that showed a d_N_/d_S_ > 1, were enriched for 11 GO Biological Process, such as response to stress (GO:0006950) and immune system process (GO:0002376), which included a number of genes involved in innate (*CCL26, CXCL13, LILRA, XCL1, PILRA* and *CMC4*) and adaptive (*MHC-B46*) immune responses. Further, in order to identify genes evolving rapidly on the capybara lineage, we compared each of 8,084 SCOs shared by capybara-guinea pig-rat. The top 5% (n = 404) of genes with high d_N_/d_S_ values for capybara-guinea pig relative to the highest d_N_/d_S_ value for guinea pig-rat orthologs were enriched for 36 GO Biological Process, including cell cycle (GO:0007049), cell death (GO:0008219), immune system process (GO:0002376), mitotic cell cycle (GO:0000278) and aging (GO:0007568).

As a complementary analysis of positive selection among genes, we used codon-based models of evolution [39] that take into account heterogeneous selection pressure at different sites within proteins, combined with a protein sequence-divergence approach (see Materials and methods). We identified a set of 4,452 SCOs using five available genomes of Hystricognath rodents: naked mole rat (*Heterocephalus glaber*), degu (*Octodon degus*), chinchilla (*Chinchilla lanigera*), guinea pig and capybara (Fig. 2). We considered genes in the top 5% of the distribution in both dimensions (i.e., the likelihood ratio tests [LRTs] of branch-site models and the protein distance index [PDI] statistic of protein p-distance) as positively selected in the capybara lineage, resulting in fifty-six candidate genes (see Materials and methods). The modest number of candidate genes is not surprising given the low levels of protein divergence between capybara and guinea pig (mean: 1%, 95% C.I.: 0.9-1.2). These 56 genes were enriched for 37 GO Biological Process, including cell death (GO:0008219), cell cycle (GO:0007049), cell proliferation (GO:0008283), growth (GO:0048590), mitotic cell cycle (GO:0000278) and immune system process (GO:0002376).

**Fig 2.**
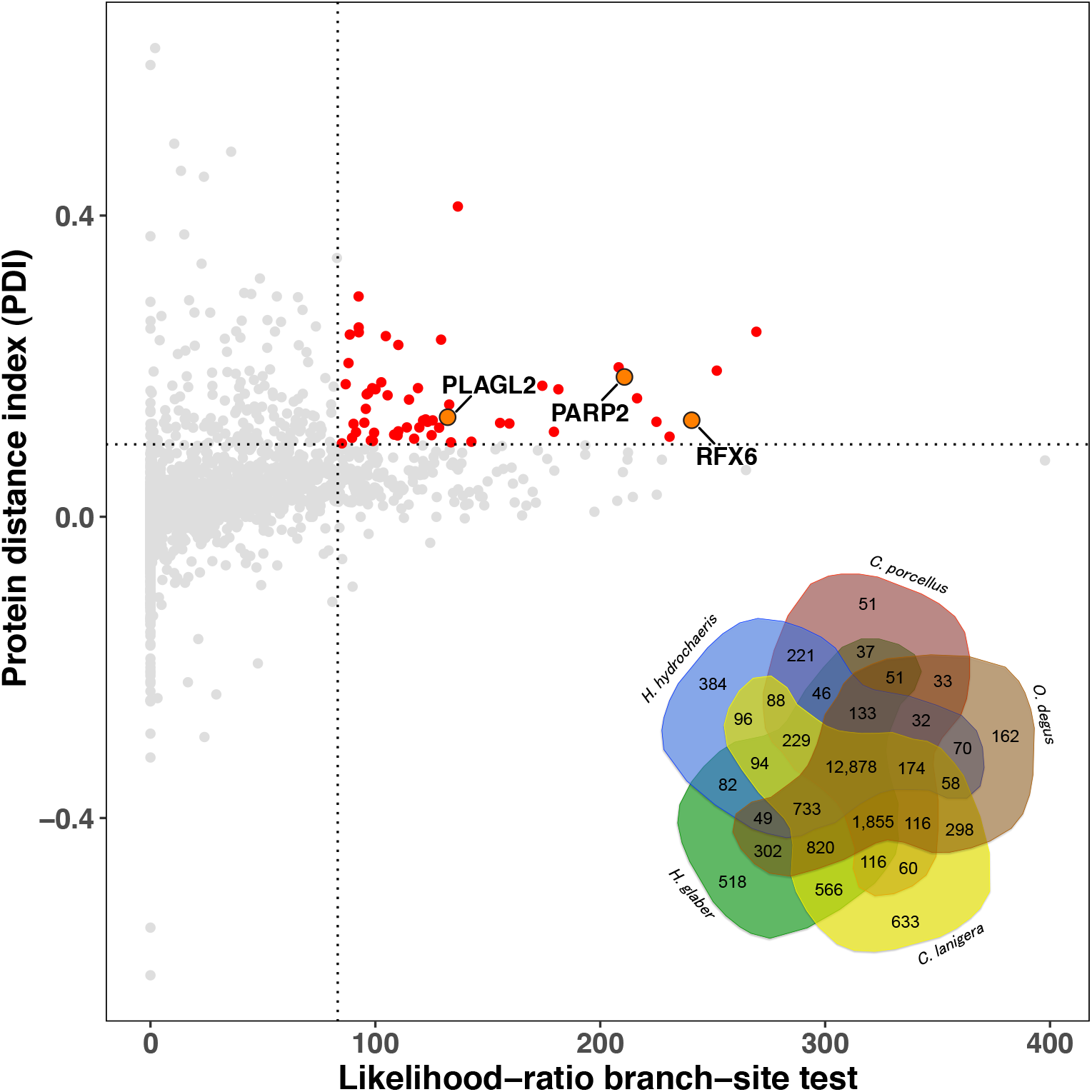
Positively selected genes in the capybara. Red dots are genes that lie in the top 5% of the distribution, above and to the right of both axes and, thus, considered to be under positive selection (see Materials and methods). Among fifty-six genes under positive selection, three were associated with growth regulation and tumor suppression (orange dots and arrows). *PLAGL2*, *Pleiomorphic adenoma-like Protein 2*; *PARP2*, *Poly(ADP-Ribose) Polymerase 2*; *RFX6*, *Regulatory Factor X6*. **Inset:** Venn diagram of the protein orthologous clusters among the 5 hystricognath species: capybara (*Hydrochoerus hydrochaeris*), guinea pig (*Cavia porcellus*), degu (*Octodon degus*), chinchilla (*Chinchilla lanigera*) and naked mole rat (*Heterocephalus glaber*). Of the 12,878 orthologous groups shared by all 5 species, 4,452 were SCOs.

### Gene family composition analysis

Gene family evolution, especially expansion by gene duplication, has been recognized as an important mechanism explaining morphological and physiological differences between species [40]. To characterize the repertoires of gene functions of rodent genomes, we employed a phenetic approach based on principal component analysis (PCA) that seeks to quantify and visualize the similarity in functional composition among species (see Matrials and methods). The PCA largely reflected phylogenetic relationships within Rodentia (Fig. 3A), except for the capybara, which showed a clear signature of genome-wide functional diversification relative to the rest of rodents. To explore which gene families were driving this pattern, we performed an overrepresentation test to determine the gene families that dominated the loadings of each principal component. The main determinants of PC1 were the diversification of gene families related to sensory perception of smell, G-protein coupled receptor signaling and immune response (Fig. 3A). Further, based on a genome-wide screen of 14,825 gene families, we identified 39 families significantly expanded in capybara, and 1 family significantly contracted.

**Fig 3.**
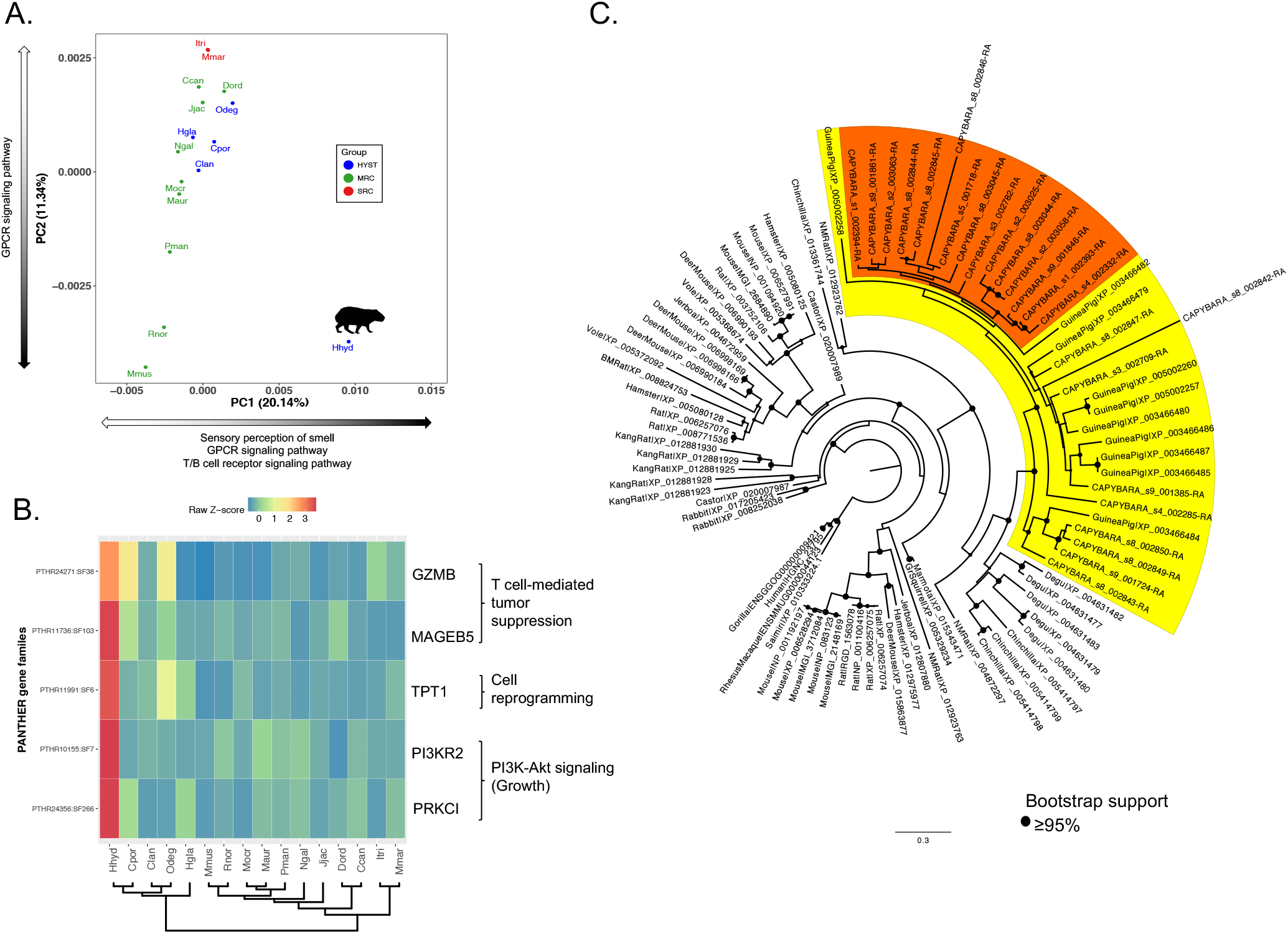
Gene family composition analysis. **(A)** Clustering of rodent genomes in a multidimensional space of molecular functions. The first two principal components are displayed, accounting for 20.14% and 11.34% of the variation, respectively. PC1 separates the capybara from the rest of the rodents based on the enrichment of gene families related to sensory perception and immune response. HYST, Hystricognathi clade; MRC, Mouse-related clade; SRC, Squirrel-related clade. Black portion of arrows indicates diversification of gene families within each PC. **(B)** Gene families enriched in the capybara. Displayed, are gene families related to growth control and tumor suppression that showed significant enrichment in the capybara (p < 0.05). The heatmap shows normalized gene counts of PANTHER molecular function categories for the 16 rodents. **(C)** Phylogenetic tree of *MAGEB5* gene family of capybara (24 MAGEB5 members) compared with rodents and rabbit; and outgroup primates (Human, gorilla, Rhesus macaque and squirrel monkey). Yellow clade contains capybara and guinea pig members; Orange clade shows a monophyletic expansion of *MAGEB5* genes in capybara. Hhyd, *Hydrochoerus hydrochaeris* (capybara); Cpor, *Cavia porcellus* (guinea pig); Clan, *Chinchilla lanigera* (chinchilla); Odeg, *Octodon degus* (degu); Hgla, *Heterocephalus glaber* (naked mole rat); Mmus, *Mus musculus* (mouse); Rnor, *Rattus norvegicus* (rat); Mocr, *Microtus ochrogaster* (vole); Maur, *Mesocricetus auratus* (hamster); Pman, *Peromyscus maniculatus* (deer mouse); Ngal, *Nannospalax galili* (blind mole rat); Jjac, *Jaculus jaculus* (jerboa); Dord, *Dipodomys ordii* (kangaroo rat); Ccan, *Castor canadensis* (beaver); Itri, *Ictidomys tridecemlineatus* (squirrel); Mmar, *Marmota marmota* (marmot).

### Evolution of regulators of growth and size

Differences in size between related organisms can reflect differences in cell number, cell size or both (Conlon and Raff, 1999). In animals, six signaling pathways regulate growth: RTK signaling via Ras, IIS via the PI3K pathway, Rheb/Tor, cytokine-JAK/STAT, Warts/Hippo, and the Myc oncogene [41]. We found several genes within the IIS (insulin/insulin-like growth factor signaling) pathway to exhibit either signatures of pairwise divergence in the capybara, such as *LEP* (*leptin*), and *PDX1* (*Pancreatic and duodenal homeobox 1*), or to be under positive selection, such as *PLAGL2* (*Pleiomorphic adenoma-like Protein 2*) and *RFX6* (*Regulatory Factor X6*; Fig. 2). Moreover, we found two gene families within the IIS pathway to be significantly expanded in the capybara relative to other rodents (Fig. 3B), namely *PI3KR2* (*Phosphatidylinositol 3-kinase Regulatory subunit β*), and *PRKCI* (*atypical protein kinase C isoform PRKC iota*).

The IIS pathway is implicated in embryonic and post-natal growth via cell proliferation [42] consistent with the possibility that large body size in the capybara has evolved through an increase in number of cells (Fig 4A). For instance, LEP is an important growth factor that regulates body weight and bone mass through the IIS pathway [43]. PLAGL2 is a member of the PLAG family of zinc finger transcription factors, which interacts with the insulin-like growth factor II (*igf2*) activating the *igf2*-mitogenic signaling pathway [44], and disruption can result in decreased body weight and post-natal growth retardation in mice [45,46]. IGF-II provides a constitutive drive for fetal and post-natal growth [47] and it has been shown that IGF-II levels do not drop after birth in caviomorph rodents, but instead remain detectable in adulthood [48], supporting a role for IGF-II regulation on the evolution of gigantism in the capybara. PI3K signalling is implicated in cell proliferation, growth, and survival via the IIS and mTOR signaling pathways [49]. PI3KR2 has been shown to have both oncogenetic [50] and tumor suppressor properties [51], and PRKCI functions downstream of PI3K and is involved in cell survival, differentiation, and proliferation by accelerating G1/S transition [52].

**Fig 4.**
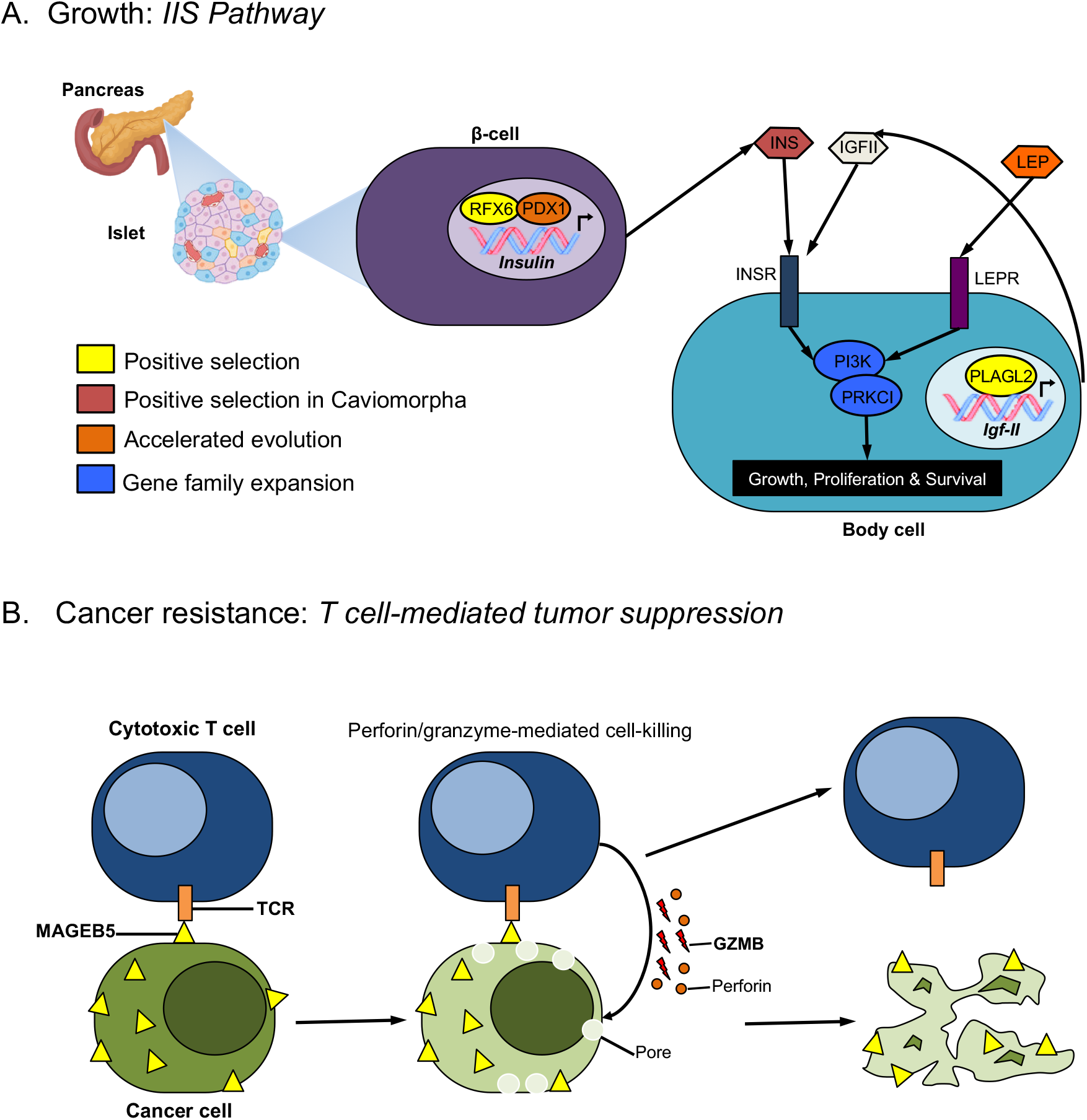
Proposed signaling pathways and mechanisms involved in (A) growth and (B) cancer resistance in the capybara. INSR: Insulin receptor; LEPR: Leptin receptor.

PDX1 and RFX6 are two key transcription factors (TFs) that regulate the development of the insulin-producing pancreatic β-cells. Pancreatic islets are composed of 5 different cell types, being the β-cells the most prominent, composing 50-80% of the islets and the ones in charge of producing insulin [53]. PDX1 and RFX6 TFs are crucial to establish and maintain the functional identity of developing and adult β-cells [54,55]. Furthermore, in adult β-cells PDX1 and RFX6 regulate insulin gene transcription [56] and, particularly, RFX6 also regulates insulin content and secretion [57]. Thus, besides the physiological effects of caviomorph insulin [26,27], lineage-specific shifts in insulin transcription and secretion profiles may be an important mechanism in the evolution of extreme body size in the capybara.

### Evolution of cancer resistance

In rodents, cancer incidence has been reported to be 46% in a wild-caught population of *Mus musculus* raised in the laboratory [58], and 14.4% to 30% in guinea pigs older than three years [59]. If capybara’s large body size is caused by an increase in cell proliferation, this could potentially lead to an increased risk of cancer. However, in capybaras, only three cases of cancer have been reported to date [60–62], suggesting the possibility of a lower incidence of cancer in capybara relative to other rodents. If so, this pattern would be consistent with empiric analyses of Peto’s Paradox [63]. Thus, we searched for genes and gene families related to cancer resistance in the capybara.

We found three gene families significantly expanded in the capybara relative to other rodents related to tumor reversion and cancer suppression by the immune system (Fig. 3B-C), namely *TPT1* (*Tumor Protein, Translationally-Controlled 1*), *MAGEB5* (*Melanoma Antigen Family B5*) and *GZMB* (*Granzyme B*). TPT1 has a major role in phenotypic reprogramming of cells, inducing cell proliferation and growth via the mTOR signaling pathway, and plays a role in tumor reversion where cancer cells lose their malignant phenotype [64]. Type I MAGE genes (e.g., MAGEB5) are expressed in highly proliferating cells such as placenta, tumors, and germ-line cells [65]. In normal somatic tissues these genes are deactivated but when cells become neoplastic, MAGE genes are reactivated and the resultant proteins may be recognized by cytotoxic T lymphocytes, triggering a T-cell-mediated tumor suppression response [65]. Lastly, GZMB is an important component of the perforin/granzyme-mediated cell-killing response of cytotoxic T lymphocytes via caspase-dependent apoptosis [66]. The fact that the capybara displays functional diversification in the T-cell receptor (TCR) signaling pathway (Fig. 3A), and expansion of MAGEB5 and GZMB gene families, suggests that T-cell-mediated tumor suppression response may be a parallel strategy, besides replicative senescence, to reduce the increased cancer risk related to the evolution of a giant body size (Fig. 4B).

Lineage-specific amino acid replacements are good predictors of change in protein function [67]. We aligned 876 SCOs from 9 rodent species spanning the three main groups within Rodentia and identified sites otherwise conserved in all rodents except for the capybara (See Materials and methods). Among the top 5% genes (n = 44) with the highest concentration of unique-capybara residues we found *TERF2-Interacting Telomere Protein 1* (*TERF2IP*). TERF2IP is part of the shelterin complex that shapes and controls the length of telomeric DNA in mammals [68], particularly the negative regulation of telomere length [69], and it has been shown that many TERF2-interacting proteins are mutated in chromosomal instability syndromes, increasing the susceptibility to cancer [70]. PARP2 (*Poly(ADP-Ribose) Polymerase* member 2), a candidate protein showing signatures of lineage-specific positive selection in the capybara (Fig. 2), directly interacts with the TERF2 protein affecting its ability to bind telomeric DNA, and disruption causes chromatid breaks that affect telomere integrity [71]. Together, these results support previous evidence showing that replicative senescence plays an important role in reducing the increased cancer risk in the big rodents [28].

### Intragenomic conflict during evolution of gigantism

How can giants evolve, or even exist? Size is regulated through somatic growth [72] and cancer appears through somatic evolution [73]. Thus, cancer places a major constraint on the evolution of large bodies. To investigate how growth and cancer suppression pathways may be coevolving to allow the evolution of a giant body size in the capybara, we used the genes that showed lineage-specific positive selection and rapid evolution, high concentration of unique-capybara residues, and capybara-expanded gene families (see above) to perform network analysis of 82 interacting genes using String Database ([74]). Network analyses based on GO Biological Process revealed six functional clusters among the 82 genes including immunity, regulation of cell proliferation, cell communication and adhesion, metabolism and homeostasis, mitochondrion organization and metabolism, and ribosome maturation (Fig. 5A).

**Fig 5.**
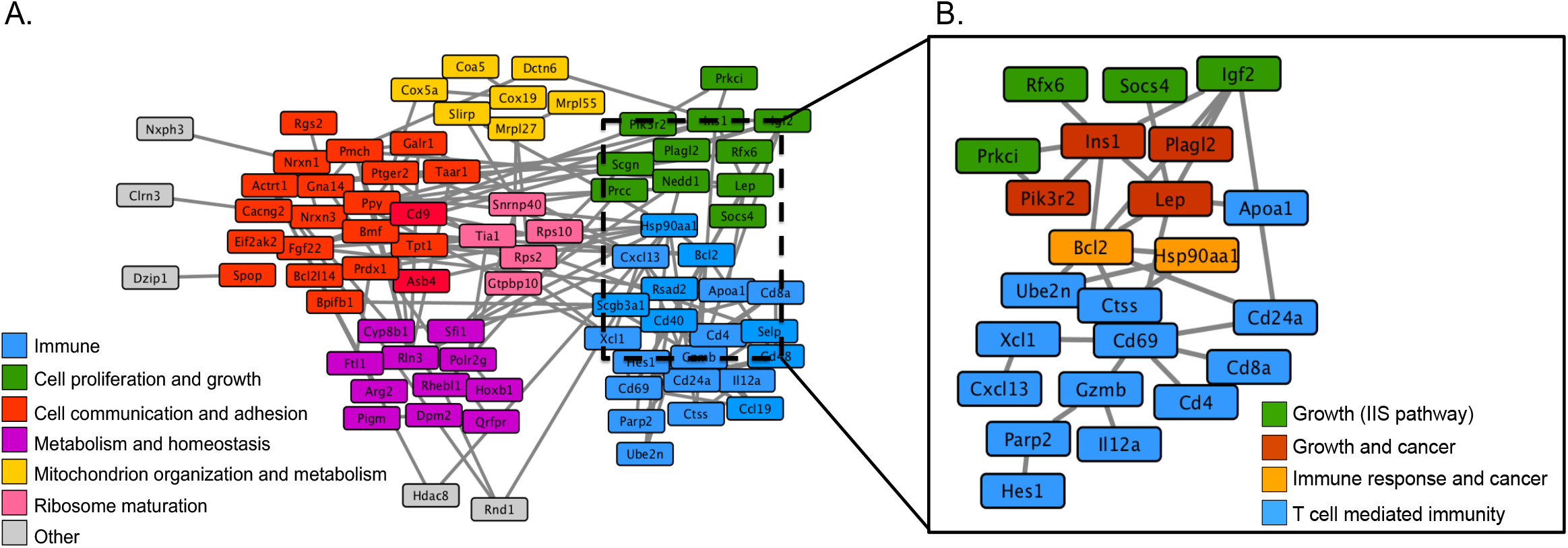
**(A) Gene interaction network of capybara genes.** Molecular interactions among 82 genes based on lineage-specific family expansions, positive selection, rapid protein evolution and unique amino acid substitutions at sites otherwise fixed in rodents. Cluster analysis was based on GO Biological Process (**B) Intragenomic conflict during evolution of gigantism.** Molecular interactions among 23 major genes involved in growth regulation by the IIS pathway, and T cell mediated immune response. 4 of the 8 genes within the IIS pathway act also as oncogenes and are involved in known cancer pathways.

We extracted 23 genes deeply involved in growth regulation by the IIS pathway, and the T cell-mediated immunity pathway to explore in more detail the interaction between growth and cancer suppression mechanisms. Remarkably, we found that this smaller network has an enrichment of cancer pathways such as prostate cancer (KEGG ID 05215), colorectal cancer (KEGG ID 05210), small cell lung cancer (KEGG ID 05222), and pathways in cancer (KEGG ID 05200). For instance, INS1, PI3KR2, PLAGL2 and LEP have been reported as growth factors, promoting somatic evolution, but also oncogenes, enabling tumor progression (Fig. 5B). Paradoxically, LEP is a major regulator of body weight and bone mass [43] but also promotes angiogenesis through VEGF signaling and acts as an autocrine factor in cancer cells promoting proliferation and inhibition of apoptosis [75]. PLAGL2 contributes to post-natal growth by activating the *igf2*-mitogenic signaling pathway [44] but also contributes to cancer by generating loss of cell-cell contact inhibition, maintaining an immature differentiation state in glioblastomas, and inducing proliferation of hematopoietic progenitors in leukemia [44,76,77]. Insulin-PI3K signaling is implicated in cell proliferation and growth and survival via the IIS and mTOR signaling pathways [49], but is frequently mutated in solid tumors [78]. These observations suggest that some cancers could arise as a by-product of growth regulatory evolution.

The dual nature of growth factors can create a scenario for an intragenomic conflict in large-sized organisms, where somatic evolution leading to gigantism may promote tumorigenic cells [79]. Cancers are selfish cell lineages that evolve through somatic mutation and cell-lineage selection [80]. Thus, selection for large body size achieved through greater cell proliferation can favor mutations that also confer cellular autonomy in growth signals enhancing their own probability of replication and transmission, a hallmark of cancer [79,81]. In fact, within species the risk of cancer scales with body size [9,82,83]. Thus, the opportunity for selfish cell lineages to arise, incidental to selection for cell proliferation, imposes a selection pressure for cancer surveillance mechanisms, generating a coupled evolution between growth-promoting and tumor-suppression pathways, a plausible underlying process causing Peto’s Paradox and a possible explanation for how giant bodies can evolve in the first place. Therefore, based on gene network interaction analyses, we propose a model for the evolution of gigantism in the capybara, and the resolution of the conflict, where adaptive evolution and duplication of genes within the IIS pathway promote somatic evolution through cell proliferation, leading to a phyletic increase in size, and the lineage-specific expansion of gene families within the T-cell mediated tumor suppression pathway as a mechanism to counteract the increased cancer risk.

### Genomic signatures of mutational load

To quantify the mutation load in capybara we used the ratio of non-synonymous substitutions to synonymous substitutions (d_N_/d_S_) as a proxy. Because most non-synonymous substitutions are deleterious or slightly deleterious [84], small populations, for which purifying selection is comparatively inefficient, tend to have elevated genome-wide fraction of fixed non-synonymous substitutions, and thus an overall higher value of d_N_/d_S_ [85]. Based on the inverse relationship between body mass and population size [8], we used body mass across 16 rodents as a proxy for *N_e_*. Bootstrap comparison of capybara-specific d_N_/d_S_ values with the rest of the rodents showed that the capybara has a higher genome-wide mean d_N_/d_S_ (0.15 vs. 0.11, respectively; two-tailed two-sample t-test, t = -103.25, *df* = 1075, *P <* 0.00001. Difference of mean d_N_/d_S_ between capybara and other rodents: 0.041; permutation test, *P* < 0.001; Fig. 6), suggesting elevated mutation load in the capybara relative to other rodents. Furthermore, analysis of body mass and nuclear d_N_/d_S_ across rodents showed that body mass explains nearly 30% of the variance in mean d_N_/d_S_ among species (R^2^ = 0.294, *df* = 14, *P <* 0.05; Phylogenetic Independent Contrasts (PIC): R^2^ = 0.332, *df* = 13, *P <* 0.05; Fig. S4A), suggesting that capybara may be part of a trend in which larger rodents experience less efficient purifying selection.

**Fig 6.**
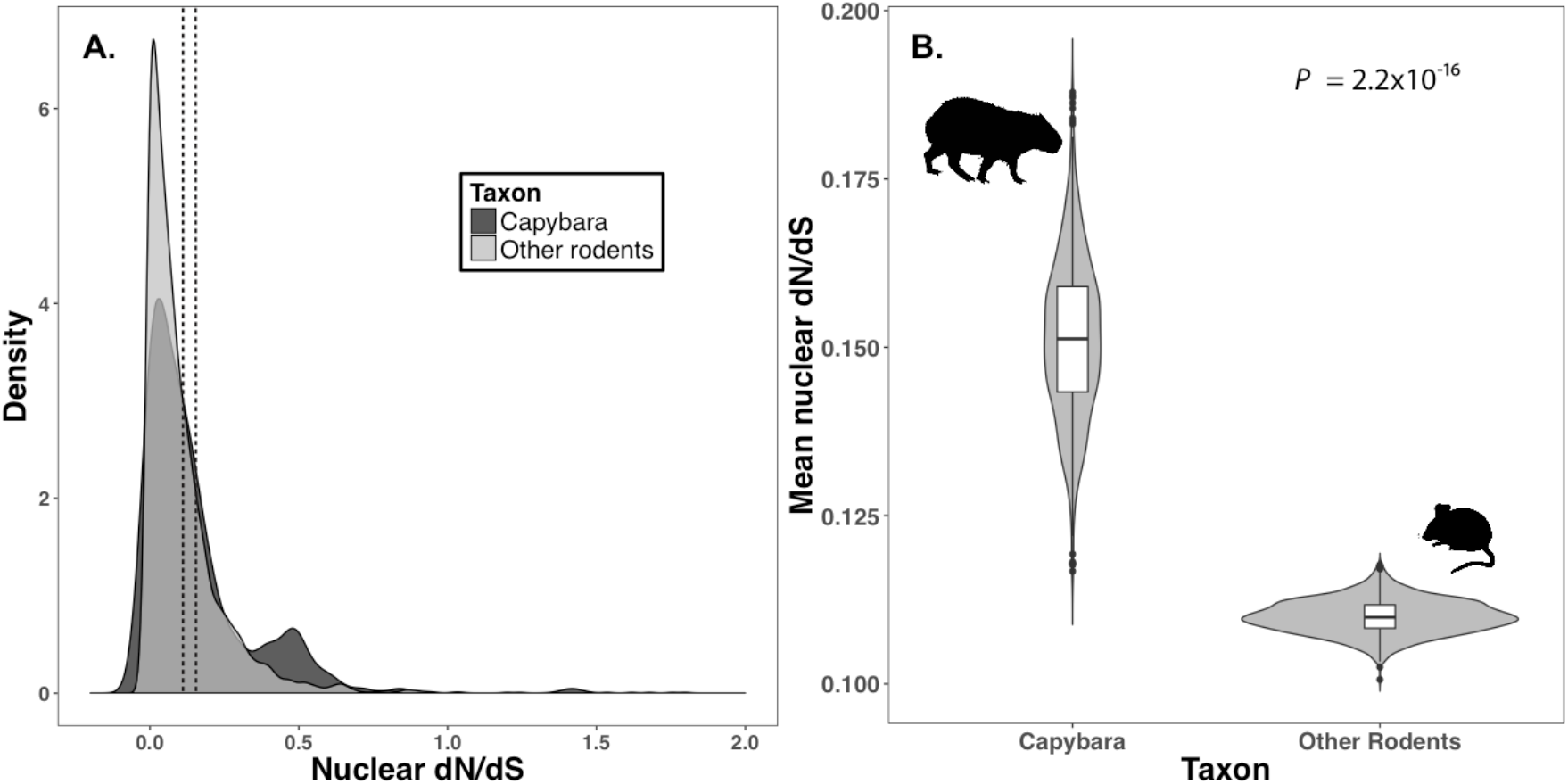
Mutation load analysis across 16 rodent genomes. The capybara shows signatures of mutational load, harboring a higher rate of accumulation of potentially deleterious mutations across 229 SCOs. **(A)** Density plot showing the distribution of d_N_/d_S_ values for the 229 SCOs for capybara and the rest of rodents. Capybaras have a slightly higher mean value of d_N_/d_S_ relative to the rest of rodents (dashed lines): 0.15 vs. 0.11, respectively. **(B)** Mean d_N_/d_S_ was significantly higher (T-test, t = -103.25, df = 1075, 95%CI = -0.042 to -0.041, *P* < 0.00001; permutation test, *P* < 0.001) for the capybara relative to the other rodents, based on 1,000 bootstrap replicates from the distributions in (A).

To further corroborate that the increased mutational load was attributable to demographic effects, we estimated d_N_/d_S_ values for the 13 protein-coding mitochondrial genes in ten rodent species (Table S3). If there is variation in the effective population size of mitochondrial DNA (mtDNA) across species, then diversity levels and substitution patterns in mtDNA should be correlated to those in nuclear DNA given that evolutionary processes that affect *N_e_* should affect both genomes [86–88]. Consistent with this idea, mean mitochondrial and nuclear d_N_/d_S_ values within rodents were strongly correlated (Spearman’s ρ = 0.67, *P <* 0.05; PIC: Spearman’s ρ = 0.7, *P <* 0.05; Fig S4B). Together, these results suggest that lower *N_e_* in larger rodent species has increased the mutation load in both nuclear and mitochondrial genomes, supporting a tradeoff at the population level of evolving a large body size.

## Discussion

Despite intrinsic constraints on the evolution of large bodies, mammalian species in multiple lineages have achieved dramatic body size changes through accelerated rates of phenotypic evolution [5,89]. The evolution of gigantism can therefore shed light on how biological systems can break underlying constraints, and mitigate or overcome the evolutionary effects of tradeoffs on whole-organism performance [90,91]. The genome of the capybara, the world’s largest rodent, is therefore crucial to addressing these questions. The genome assembly analyzed here has revealed insights into the capybara’s particular morphology, offering the first genomic scan onto the developmental regulators of its large size, as well as the characterization of a putative novel anti-cancer mechanism, and the collateral effects at the population level of evolving a large body size. For instance, reduced long-term *N_e_* can increase the probability of extinction because of the accumulation of slightly deleterious mutations that reduce the mean fitness of the population [12,92,93]. This may explain macroevolutionary patterns of extinction risk related to body size [94], reinforcing the effect of body size at the population level and its consequences to genome evolution.

Changes underlying body size evolution affect other life-history traits, such as fecundity and longevity [2], thus, the extent of pleiotropic effects of each developmental regulator of body size will ultimately determine its availability for evolutionary change [95]. The IIS pathway has also been linked to rapid and extreme body-size evolution in the sunfish (*Mola mola*), the world’s largest bony fish [96], and to extreme size differences in dog breeds [97]. Given the existence of at least six size-regulating pathways [41], the repeated use of the IIS pathway in fishes, dogs and rodents, suggests that this pathway might be less constrained and could be a ‘hot-spot’ for extreme body size evolution in vertebrates, indicating a common genetic and developmental pathway controlling rapid body size evolution through cell proliferation.

Finally, cancer imposes a strong constraint on morphological and ontogenetic evolution [90], particularly during the evolution of gigantism. The evolution of compensatory cancer-protective mechanisms, thus, becomes a requirement if animals are to evolve larger bodies. In fact, larger and long-lived mammals have evolved enhanced mechanisms of cancer suppression as revealed by the lower risk of developing cancer relative to smaller species, a phenomenon known as Peto’s Paradox. While some tumor suppression pathways are conserved among animals, some are the result of idiosyncratic evolutionary processes within species, resulting in a high variety of lineage-specific solutions [98]. For instance, elephants, whales and capybaras, which have evolved extremely large sizes, each have a specific solution to the inherent higher risk of cancer: higher sensitivity to DNA damage, more efficient DNA repair, and enhanced immune response against tumors, respectively. Mammalian comparative genomics to study cancer can therefore provide insight into the common processes as well as the specific mechanisms of cancer resistance, and offer potential targets for human cancer prevention and treatment. Thus, given the increasing interest in using genetically engineered T cells for cancer immunotherapy [99], the capybara could be a suitable model to study the evolution and physiology of tumor detection and eradication by T cells.

## Materials and methods

### Supplemental materials and data availability

Genome assembly: NCBI BioProject: PRJNA399400; NCBI assembly accession number: PVLA00000000.

Code with statistical analyses and tables with genes: https://github.com/santiagoha18/Capybara_gigantism

### Ethics statement

All procedures involving live animals followed protocols approved by the *Comité Institucional de Uso y Cuidado de Animales de Laboratorio* (CICUAL) of the Universidad de los Andes, approval number C.FUA_14-023, 5 June 2014. Research and field collection of samples was authorized by the *Autoridad Nacional de Licencias Ambientales* (ANLA) *de Colombia* under the *permiso marco resolución No.* 1177 to the Universidad de los Andes.

### Rate of Body Size Evolution and Ancestral State Reconstruction

The term ‘gigantism’ is relative to the clade of interest. We used a comparative approach to test whether the capybara’s large size is due to lineage-specific accelerated rates of phenotypic evolution. From the literature we obtained a recent time-calibrated phylogeny of Caviomorpha [100] and for each species obtained mean adult body mass (in grams) as the mean of the two sexes [101,102]. Analyses were conducted on natural log-transformed body mass.

The Brownian motion (BM) model of continuous trait evolution assumes that trait variation accumulates proportionally through time, rates of trait evolution are finite and constant across the phylogeny, where branch lengths are scaled to time [103]. The disparity in body size among caviomorph rodents (Fig S5), suggests that rate of body size evolution likely varies across time or among lineages. To compare the BM model with a multiple rate model of body size evolution on the caviomorph phylogeny, and to reconstruct ancestral states, we used the software STABLETRAITS [104]. STABLETRAITS samples from a symmetrical distribution of rates with mean zero and defined by its index of stability (α). BM α=2 implies a normal distribution (*i.e*., single-rate model), whereas α<2 results in a shallower platykurtic distribution with heavy tails, supporting a multi-rate model. For the analysis, results from a multi-rate model (α<2) were compared with a BM model (α=2) to select the best fitting model based on the Deviance Information Criterion (DIC). Two MCMC chains were run for 2 million generations each until the potential scale reduction factor went bellow 1.01, which implies convergence between the two chains. Burn-in was set to 25% of samples, with the output containing calculated rates, ancestral states, and posterior probabilities.

Given that the multi-rate model was selected as the best model according to the DIC (Table 1), the R package AUTEUR [34] was used to identify which branches on the phylogeny experienced rate shifts. AUTEUR uses a reversible-jump MCMC (rjMCMC) to assess the fit of models that differ in the number of rate shifts in the tree. The algorithm assumes that evolutionary rates are inherited from successive ancestral nodes, unless the data provide evidence of a rate shift. The rjMCMC approach explores models ranging from a single-rate BM process to a maximally complex process where each branch evolves under a different rate. Two rjMCMC chains were run for 1 million generations each with the following parameters: sample.freq = 100 prop.mergesplit = 0.1 and prop.width = 0.1.

### Genome size estimation

Genome size in the capybara was estimated using flow cytometry on blood samples from two juveniles taken from a population in Cabuyaro, Meta, Colombia (04.284025°, -072.725647°; Table S2). Animals where tranquilized with an intramuscular dose of Zoletil® 50 (Virbac^TM^, Colombia), and then sacrificed with a lethal dose of Euthanex® 50 (Invet^TM^, Colombia) according to manufacturer instructions, and different tissues were collected for future RNA-seq work. Blood samples were stored in 3mL heparinized tubes at 2ºC and transported to the laboratory in Bogotá, Colombia. Nucleated white blood cells were separated from the rest of blood contents by centrifugation. 100µL of white blood cells were washed with 3mL of PBS buffer, and DNA was stained according to the BD Cycletest^TM^ Plus DNA kit instructions (BD Biosciences^TM^). Chicken red blood cells were used as a genome size standard, assuming a haploid C-value of 1.25 pg. Flow cytometry was conducted on a BD FACSCanto^TM^ II cytometer with a FACs DIVA v6.1 software. Average C-value was estimated to be 3.21 pg (ANDES-M 2304 C-value = 3.17 pg, ANDES-M 2303 C-value = 3.25 pg). Converting the mean C-value to numbers of base pairs we obtained an estimated genome size of 3.14 Gb.

### Capybara genome sequencing and assembly

As part of the 200 Mammalian Genomes project, the SciLifeLab at Uppsala University and the Broad Institute sequenced and assembled a DISCOVAR *de novo* genome assembly [105] from a sample of a wild-born female *H. hydrochaeris* from the San Diego Zoo’s Frozen Zoo (frozen sample KB10393), imported from Paraguay. The genome assembly was performed with the DISCOVAR *de novo* pipeline, generating an assembly spanning 2.734 gigabases (Gb) with a scaffold N50 length of 0.202 megabases (Mb; Table 1).

The Dovetail Genomics^TM^ proprietary scaffolding method based on Chicago^TM^ libraries [106] was used to upgrade the initial DISCOVAR assembly. Dovetail’s novel approach to increasing the contiguity of genome assemblies combines initial short-read assemblies with long-range information generated by *in vitro* proximity ligation of DNA in chromatin. From the same individual of *H. hydrochaeris* used to obtain the initial DISCOVAR *de novo* assembly, 500 ng of high molecular weight (HMW) gDNA (50 Kb mean fragment size) was used as input for the Chicago^TM^ libraries. The libraries were sequenced on two lanes of an Illumina HiSeq in Rapid Run Mode to produce 316 million 2×100 bp paired-end reads, which provided 67X physical coverage (measured in bins of 1-50 kb).

The reads were assembled by Dovetail Genomics^TM^ using the HiRise^TM^ Scaffolder pipeline and the DISCOVAR *de novo* genome assembly as inputs. Shotgun and Chicago^TM^ library sequences were aligned to the draft input assembly using a modified SNAP read mapper (http://snap.cs.berkeley.edu) to generate an assembly spanning 2.737 Gb, with a contig N50 length of 161.1 kb and an impressive scaffold N50 length of 12.2 Mb, as calculated using the QUAST software [107].

### Assembly quality and annotation

Two approaches were employed to evaluate the quality of the assembly: 1) the Core Eukaryotic Genes Mapping Approach [31] was used to determine the number of complete Core Eukaryotic Genes (CEGs) recovered, and 2) an analysis of Benchmarking Universal Single-Copy Orthologs [30] was applied as an evolutionary measure of genome completeness, using the vertebrate dataset (3023 genes) as query. Gene content of a well-assembled genome should include a high proportion (≥75%) of both CEGs and BUSCOs (Fig S1).

Putative genes were located in the assembly by homology-based annotation with MAKER v2.31.9 [32] based on protein evidence from mouse (*Mus musculus*), rat (*Rattus norvegicus*) and guinea pig (*Cavia porcellus*; Ensembl release 85). CD-HIT v4.6.1 [108] was used to cluster highly homologous protein sequences across the three protein sets and generate a non-redundant protein database (nrPD), which was used to guide gene predictions on the capybara genome. To evaluate the quality of the annotations, the Annotation Edit Distance (AED) score was used as a measure of congruence between each annotation and the homology-based evidence (in this case, protein sequences). When an annotation perfectly matches the overlapping model protein, the AED value is 0 [109]. As a rule of thumb, a genome annotation where 90% of the annotations have an AED less than 0.5 is considered well annotated (Campbell et al., 2014). The capybara genome annotation has 92.5% of its annotations with an AED below 0.5.

PANTHER [110] was used to characterize the functions of the capybara-annotated proteins (Fig S2). PANTHER is a library of protein families and subfamilies that uses the Gene Ontology (GO) database to assign function to proteins. Following the protocol for large-scale gene function analysis with PANTHER [111], the Scoring Tool was used to assign each protein of the capybara genome to a specific protein family (and subfamily) based on a library of Hidden Markov Models.

### Annotation of repeat elements

RepeatMasker open-4.0.5 [112] was used to identify and classify transposable elements (TEs) and short tandem repeats by aligning the capybara genome sequences against a reference library of known repeats for rodents, using default parameters. For comparison, NCBI Annotation releases for 13 additional rodents were used (Fig. 1)

### Phylogenomics and divergence time estimation

Available glire (rodent + rabbit) genomes were downloaded from NCBI and Ensembl (Table S3). AlignWise [113] was used to identify protein-coding regions and extract the longest open reading frame per gene. Single-copy gene orthologs (SCOs) shared across the 17 glire genomes (Table S3) were identified using an 8321-marker dataset of orthologs for glires from OrthoMaM database v8 [114]. This dataset contains orthologous groups for 6 of the 17 genomes; thus, we constructed a dataset combining the 8321 OrthoMaM orthologous groups with the 11 remaining genomes and performed an all-vsall blastn [115]. We used the *reciprocal groups* function of VESPA v1.0β [116] to recover orthologous groups spanning the 17 genomes and identified 2597 groups, of which 370 were SCOs. For phylogenetic tree inference, each of the 370 SCOs was aligned using MUSCLE [117] and then concatenated. Poorly aligned sites were trimmed using Gblocks v0.91b using the –t=c option to retain the reading frames [118]. RAxML v8.2.10 [119] with rapid bootstrap mode (n = 500) and assuming a GTRGAMMA model was used to infer a phylogenetic tree. We also used a species-tree approach to phylogenomic inference that takes into account the individual history of each gene, which may conflict due to, e.g., incomplete lineage sorting, using the software Neighbor-joining species tree (NJst; [120]). For each individually aligned SCO, 100 bootstrap replicates were run on RAxML v8.2.10. A file with 37000 gene trees (100 bootstraps x 370 SCOs) was used as input for NJst.

Given that both phylogenomic analyses (concatenated and species-tree approach) resulted in identical topologies, divergence times were estimated based on the topology obtained in the concatenated analysis using the software MCMCTREE [121] in the PAML package v4.8 [39]. Two nodes were calibrated using fossil data [122] as follows: the most recent common ancestor (MRCA) of Rodentia was restricted to the interval 76 – 88 million years ago (Ma), and the MRCA of *Mus* and *Rattus* was limited to 18 – 23 Ma.

### Genomic signatures of mutation load

In order estimate the mutation load, we used the ratio of the rates of non-synonymous substitutions over synonymous substitutions (d_N_/d_S_) as a proxy. A set of 229 SCOs shared across the 16 rodent genomes (Table S3) was recovered with the software Proteinortho [123]. Values of d_N_/d_S_ were estimated for the set of 229 SCOs, using the program CODEML from the PAML package v4.8 [39]. Likewise, d_N_/d_S_ ratios for the 13 protein-coding genes of the mitochondrial genomes of capybara plus 9 additional rodent mitochondrial genomes were estimated (Table S3). Two evolutionary models were evaluated: model 0 with d_N_/d_S_ held constant over the time-calibrated concatenated likelihood tree, and model 1 with free d_N_/d_S_ estimated separately for each branch. Likelihood-ratio tests were used to compare the two models, and in each case the model 1 was the preferred one. Only d_N_/d_S_ values associated with external branches were used as measures of mutation load. Codon-based alignments were performed with MACSE [124] with default parameters and poorly aligned regions were trimmed with Gblocks v. 0.91b [118].

To determine if the capybara has a higher genome-wide d_N_/d_S_ relative to other rodents, we (a) calculated the mean ratio from each of 1000 bootstrapped samples of the original 229 capybara-specific d_N_/d_S_ values (see above), and then did the same for the combined set of d_N_/d_S_ values for the other 15 rodent genomes. The two samples of 1,000 means were then compared using a two-sample *t*-test. (b) A permutation test (1,000 permutations) was performed to create a null distribution of the differences of (mean capybara d_N_/d_S_ – mean rodents d_N_/d_S_), and then compared to the observed value: 0.041.

To test whether variation in d_N_/d_S_ was caused by demographic effects (i.e. higher genetic drift relative to purifying selection), the correlation between nuclear and mitochondrial d_N_/d_S_ values across species was estimated, under the hypothesis that a demographic bottleneck should affect both genomes [87]. The correlations between (a) Mean nuclear d_N_/d_S_ versus body mass and (b) mean nuclear d_N_/d_S_ versus mean mitochondrial d_N_/d_S_, were re-evaluated using phylogenetic independent contrasts (PICs; [125]) to control for the potentially confounding effects of shared evolutionary history. Because for most mammalian species comparable genetic estimates of nuclear *N*_e_ were not available, we used body mass as a proxy for *N*_e_. Mean d_N_/d_S_ values per species and natural log-transformed values of body mass were used to estimate the impact of the reduction of effective population size on the strength of purifying selection. Assessment of linear model assumptions was performed with the gvlma R package [126].

### Gene family composition

To search for expanded gene families in the capybara, we performed gene family assignments for each of the 16 rodent genomes (Table S3) using the protocol for large-scale gene function analysis with PANTHER [111]. Following [127], functional repertoires of the genomes were represented in a multidimensional space, where each dimension corresponds to a particular gene family. A principal component analysis (PCA) was conducted using the prcomp function in R [126] on the count of the number of genes in each gene family as identified by PANTHER. The coordinate of a species’ genome along each dimension represents the number of genes that it contains with the corresponding gene family normalized by the total number of genes in that particular species. PCA based on content and size of gene families has been shown to reflect important evolutionary splits between groups or clades of animals (non-bilaterian and bilaterian: [127]; deuterostomes and protostomes: [128]). In rodents, PC1 component separated the capybara from all other rodent genomes and explained the 20.14% of variance, while PC2 grouped all genomes and explained 11.34% of the variance. An overrepresentation test was conducted on the gene families that fell on both 2.5%-tails of the loadings of PC1 and PC2, to determine the gene families that dominate the loadings of each principal component. A Fisher’s exact test was then performed iteratively in R (R Development Core Team, 2008) on counts of each PANTHER gene family from the 5% tails of PC1 and PC2 against the complete set of gene families (23,110 families). To be defined as enriched, we assumed a significance level α = 0.05 for each gene family, before a traditional Bonferroni correction for multiple testing.

To conduct a genome-wide screen for gene family expansions, a table with 14,825 shared PANTHER gene families, present in at least 15 of the 16 genomes, was constructed with the counts for genes annotated to those families in the 16 rodent genomes (see above). A Fisher’s exact test was then performed iteratively in R [126] comparing the number of genes found in the capybara in each gene family against the total number of genes in the remaining 15 rodents for a given gene family. To be defined as enriched, a gene family had to have a significant p-value (α<0.05) after a traditional Bonferroni correction for multiple testing. We calculated a Z-score per gene family per species, where 1) the individual counts of each gene family per species were normalized by the total number of genes in the genome of each species and 2) was standarized within each gene family (across species) with the scale function in R [126] and visualized as a heatmap (Fig. 7B). This Z-score represented the number of standard deviations below or above the mean gene family count for each species.

### Pairwise d_N_/d_S_ analysis

Pairwise comparisons of guinea pig and capybara could provide insights into the genes underlying capybara’s large size, as well as insights into other adaptive differences. We calculated pairwise d_N_/d_S_ ratios of each gene within genomes to look for signs of positive selection. A total of 11,255 single-copy gene orthologs (SCOs) between guinea pig and capybara where obtained with the OrthoFinder software v.1.1.4 [129]. Specific Biopython [130] and PyCogent [131] modules where used to estimate d_N_/d_S_ values for each SCO. To identify genes that evolved more rapidly in the capybara lineage, we calculated the ratios of the capybara-guinea pig d_N_/d_S_ values relative to the highest d_N_/d_S_ value for guinea pig-rat, on a set of 8,084 SCOs, and ranked them by decreasing ratio. Codon-based alignments were performed with MACSE [124] with default parameters. Gaps and ambiguities were excluded from alignments, and only alignments with a trimmed length above 50% of the original alignment length were retained for subsequent analyses.

### Positive selection analysis using codon-based models of evolution

While pairwise d_N_/d_S_ can identify genes with overall faster sequence evolution, amino acid sites within a protein may experience different selective pressures and have different underlying d_N_/d_S_ ratios [132]. We fitted branch-site models to identify positive selection in a set of 4,452 SCOs obtained using the OrthoVenn software [133] on capybara versus four closely related rodents with published whole genome sequences: naked mole rat, degu, chinchilla and guinea pig. Downstream selective pressure variation analyses were done using the Selection Analysis Preparation functions provided in VESPA v1.0β [116]. We compared the likelihood scores of two selection models implemented in CODEML in the PAML package v4.8 [39] using likelihood-ratio tests (LRT) as a measure of model fit: a branch-site model (model A) in which attempts to detect positive selection acting on a few sites on specified lineages or ‘foreground branches’ (d_N_/d_S_>1), versus the null model (A null) in which codons can only evolve neutrally (d_N_/d_S_ =1) or under purifying selection (0< d_N_/d_S_ <1; [134]). As maximum likelihood methods to detect positive selection are sensitive to sample size [135], increasing the possibility of false positives [136], we calculated a second measure of rapid evolution of each gene: ((Capybara, guinea pig), degu) protein trios where used to estimate the difference of the raw p-distance in protein sequence between capybara and degu (pdCD), and the raw p-distance between guinea pig and degu (pdGD). These two distances were used to calculate the ‘protein distance index’ or PDI for each *i* gene (PDI_*i*_ = pdCD_*i*_ – pdGD_*i*_) which takes a value of zero if the two p-distances are equal, meaning that the capybara and guinea pig lineages have evolved at equal rates since their divergence from their MRCA. A positive value is indicative of accelerated protein evolution in the capybara lineage. We considered genes in the top 5% of the distribution in both dimensions (i.e., the LRTs of branch-site models and the PDI statistic) as positively selected in the capybara lineage. Codon-based alignments were performed with MACSE [124] with default parameters and poorly aligned regions were trimmed with Gblocks v. 0.91b [118].

For genes exhibiting signatures of lineage-specific positive selection, we conducted GO enrichment analysis to identify Biological Process overrepresented in these genes relative to the 4,452 SCO set. The GO Biological Process for each gene was retrieved with the biomaRt R package [137] using the protein stable IDs of guinea pig orthologs. A Fisher’s exact test was then performed iteratively in R [126] on counts of each Biological Process from the positively selected genes against the full set of orthologs (4,452 genes). To be defined as enriched, a GO Biological Process had to have a significant p-value (α<0.05) after a traditional Bonferroni correction for multiple testing.

### Identification of proteins with capybara-unique residues

Conservation-based methods used to identify lineage-specific amino acid replacements have been proven to be good predictors of change in protein function [67]. We identified SCOs for 9 rodents (capybara, guinea pig, degu, chinchilla, naked mole rat, Ord’s kangaroo rat, mouse, rat and thirteen-lined ground squirrel) and used multiple sequence alignments (MSA) to identify positions in each SCO that are conserved in all species except for the capybara. An in-house Python script was used to identify unique capybara amino acid residues where a maximum of one unknown residue was allowed in each position of the alignment. SCOs were identified with OrthoFinder software v.1.1.4 [129] and protein MSAs were performed with PRANK [138] with default parameters. Gaps were excluded from alignments, and only alignments with a trimmed length above 50% of the original alignment length were retained for subsequent analyses. This procedure resulted in a total of 876 SCOs retained. The SCOs were ranked according to the number of unique substitutions normalized by protein length.

### Gene interaction network analyses

We generated a list of 195 candidate genes that showed lineage-specific positive selection, pairwise dN/dS > 1, rapid evolution (top 1%), high concentration of capybara-unique residues (top 5%) and gene-family expansions specific to the capybara. Insulin was included as part of the 195 gene set given that previous analyses showed signatures of adaptive evolution among Caviomorph rodents [26]. Of the 195 genes, only 82 genes presented known or predicted interactions with each other according to String Database [74]. We used this new gene set to perform a network analysis of GO Biological Process and KEGG pathways on the 82 interacting genes. Networks were constructed based on String DB interactions and plotted with Cytoscape 3.0 [139]. GO and KEGG enrichment analyses were performed based on String DB information with a false discovery rate of 0.05.

## Acknowledgements

This work was supported by *Colciencias* grant in Science, Technology & Innovation BIO number 1204-659-44334 (A.J.C.); seed grant from the *Facultad de Ciencias*, Universidad de los Andes (S.H.A., A.J.C.). Genome sequencing and assembly was supported by the 200 Mammals Project at the Broad Institute (NIH Grant 5R01HG008742-02) and Uppsala University (Swedish Research Council). Special thanks to: Diane Genereux, Daniel Cadena and Jeremy Johnson for insightful comments on the final version of the manuscript; Eva Murén and Voichita Marinescu for generating the sequencing libraries; Juanita Herrera and veterinarian Marlly Guarín for help with drafting and implementing animal care and use protocols; Universidad de los Andes’ HPC group for cluster access; John Marío González in the medical school of the Universidad de los Andes for access and help with flow cytometry; *Agencia de Licencias Ambientales* (ANLA) for collecting permits granted to Universidad de los Andes (permit Nº 1177).

## Author contributions

A.J.C., E.K and K.L.T. conceived genome-sequencing project. O.A.R. provided materials. E.K. and K.L.T. provided the DISCOVAR genome. A.J.C. provided financing. S.H.A. conceived the evolutionary questions addressed here, performed all evolutionary and statistical analyses, and wrote the manuscript. All authors revised and approved final version.

## Figures and tables

**Table S1.**
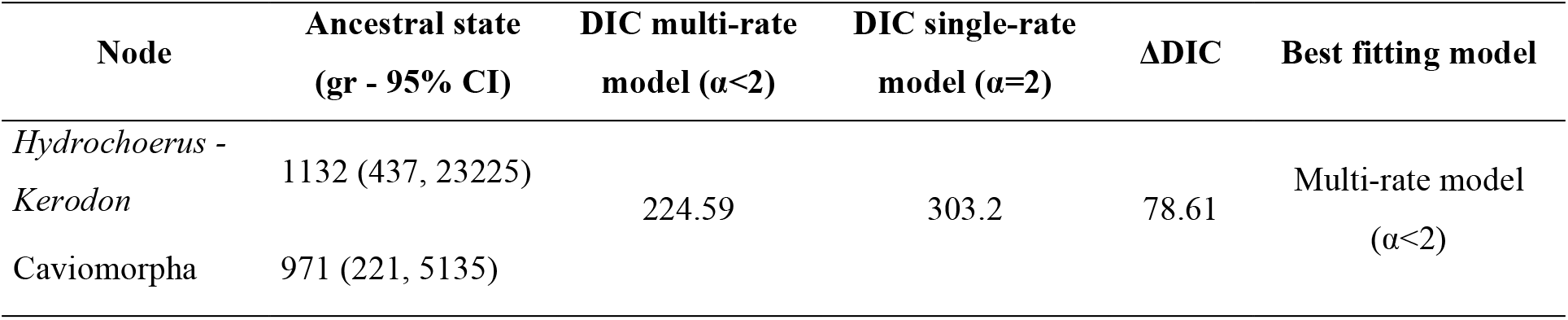
Reconstruction of ancestral body mass for the caviomorph clade showed that capybara likely evolved from a relatively small ancestor. Body mass reconstruction was done with STABLETRAITS [104], which accounts for rate variation in trait evolution along a phylogeny. Results from a multi-rate model (α<2) were compared with a BM model (α=2) to select the best fitting model based on the Deviance Information Criterion (DIC).

| Node | Ancestral state<br>(gr - 95% CI) | DIC multi-rate<br>model ($\alpha < 2$) | DIC single-rate<br>model ($\alpha = 2$) | ΔDIC | Best fitting model |
| --- | --- | --- | --- | --- | --- |
| *Hydrochoerus* -<br>*Kerodon* | 1132 (437, 23225) | 224.59 | 303.2 | 78.61 | Multi-rate model<br>($\alpha < 2$) |
| Caviomorpha | 971 (221, 5135) |  |  |  |  |

**Table S2.**
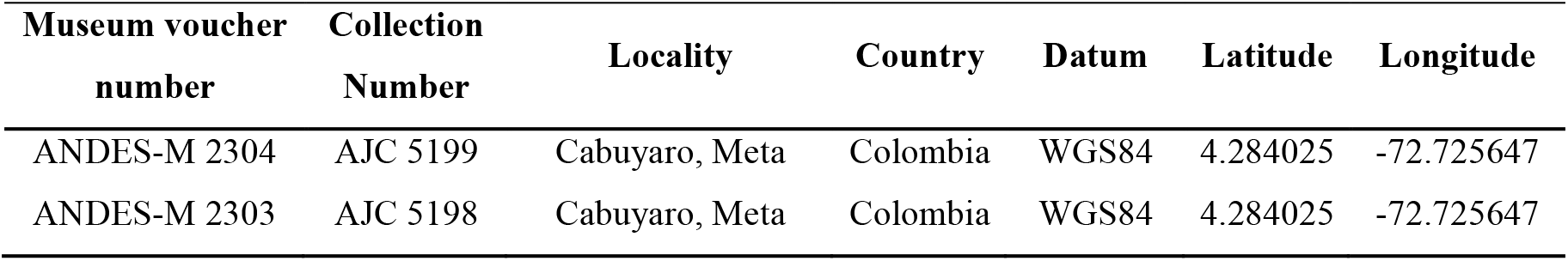
Field and collection information of the two specimens analyzed for genome size estimation.

**Table S3.** Published genomes used for comparative analyses. ^a^ Species used for mitochondrial genome analyses. Mitochondrial genomes were downloaded from the Organelle Genome Resources database of NCBI. ^b^ Direct submission to NCBI (Unpublished data).

| Clade | Species | Genome version | Database | Reference |
| --- | --- | --- | --- | --- |
| Hystricognathi | Guinea pig (*Cavia porcellus*)<sup>a</sup> | cavPor3 | ENSEMBL | [140] |
| Hystricognathi | Long-tailed chinchilla (*Chinchilla lanigera*)<sup>a</sup> | ChiLan1.0 | NCBI | NCBI<sup>b</sup> |
| Hystricognathi | Degu (*Octodon degus*)<sup>a</sup> | OctDeg1.0 | NCBI | NCBI<sup>b</sup> |
| Hystricognathi | Naked mole rat (*Heterocephalus glaber*)<sup>a</sup> | HetGla_female_1.0 | NCBI | [141] |
| Mouse-related clade | Mouse (*Mus musculus*)<sup>a</sup> | GRCm38.p5 | ENSEMBL | [33] |
| Mouse-related clade | Rat (*Rattus norvegicus*)<sup>a</sup> | Rnor_6.0 | ENSEMBL | [142] |
| Mouse-related clade | Prairie vole (*Microtus ochrogaster*) | MicOch1.0 | NCBI | NCBI<sup>b</sup> |
| Mouse-related clade | Golden hamster (*Mesocricetus auratus*)<sup>a</sup> | MesAur1.0 | NCBI | NCBI<sup>b</sup> |
| Mouse-related clade | North American deer mouse (*Peromyscus maniculatus*) | Pman1.0 | NCBI | NCBI<sup>b</sup> |
| Mouse-related clade | Blind mole rat (*Spalax galili*)<sup>a</sup> | S.galili_v1.0 | NCBI | [143] |
| Mouse-related clade | Lesser Egyptian jerboa (*Jaculus jaculus*)<sup>a</sup> | JacJac1.0 | NCBI | NCBI<sup>b</sup> |
| Mouse-related clade | Ord's kangaroo rat (*Dipodomys ordii*) | Dord_2.0 | NCBI | [140] |
| Mouse-related clade | North American beaver (*Castor canadensis*) | C.can genome v1.0 | NCBI | [144] |
| Squirrel-related-clade | Thirteen-lined ground squirrel (*Ictidomys tridecemlineatus*) | SpeTri2.0 | NCBI | [140] |
| Squirrel-related-clade | Alpine marmot (*Marmota marmota*) | marMar2.1 | NCBI | NCBI<sup>b</sup> |
| Outgroup | Rabbit (*Oryctolagus cuniculus*) | OryCun2.0 | NCBI | [140] |

**Fig S1.**
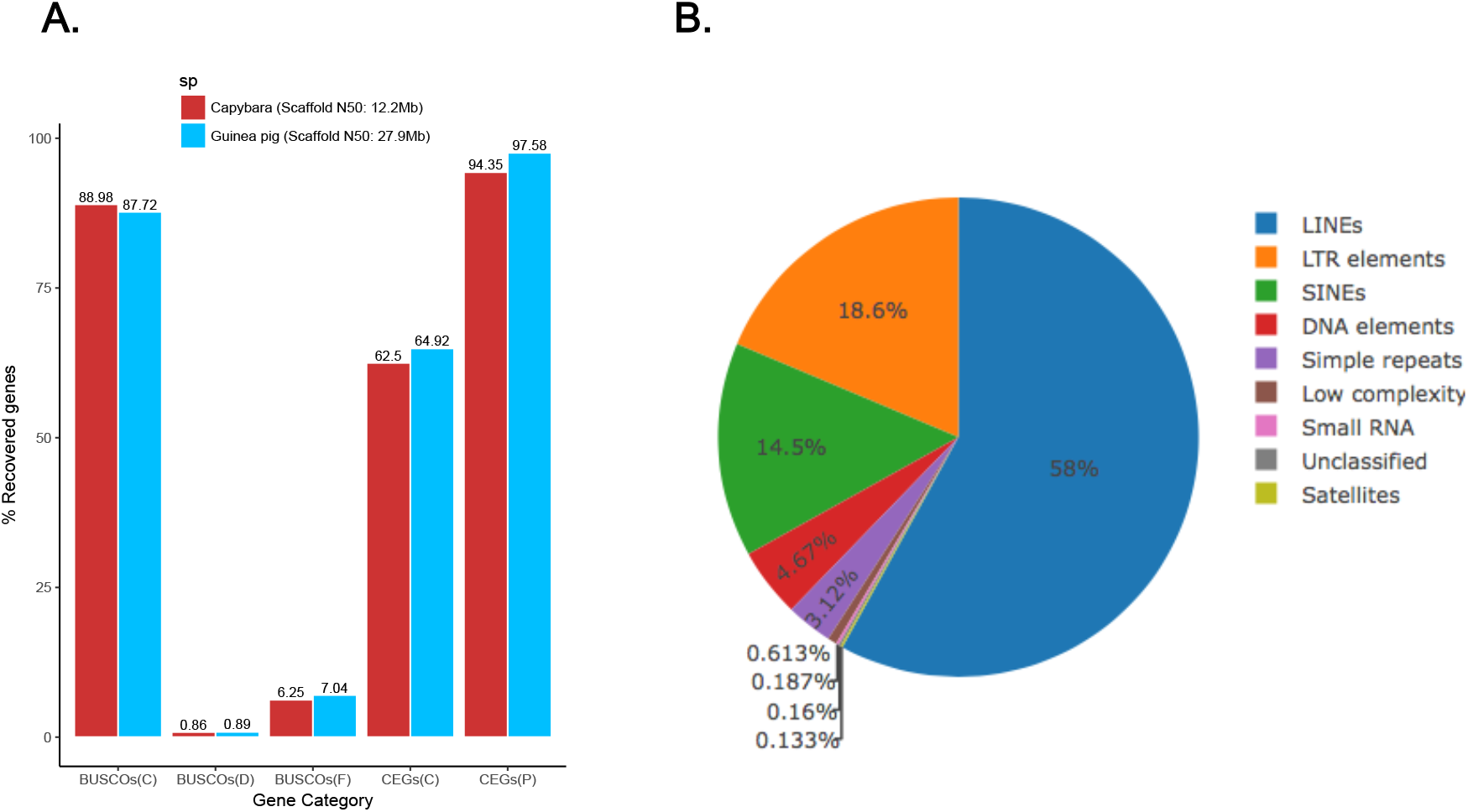
Capybara genome annotation statistics. **(A)** Quality assessment of the capybara genome compared to the guinea pig genome. CEGMA analysis was conducted for 248 CEGs. BUSCO analysis was conducted with the vertebrate dataset that included 3023 BUSCOs. Note that guinea pig scaffold N50 is twice as high as the capybara scaffold N50, but both genomes are of similar quality. (C): Complete, (D): Duplicated, (F): Fragmented, (P): Partial. **(B)** Estimated proportion of the capybara genome sequence occupied by repetitive elements. Results are based on identification of known and *de novo* repeat elements aligning the capybara genome sequences against a reference library of known repeats for rodents using RepeatMasker. LINE, long interspersed element; LRT, long terminal repeat; SINE, short interspersed element.

**Fig S2.**
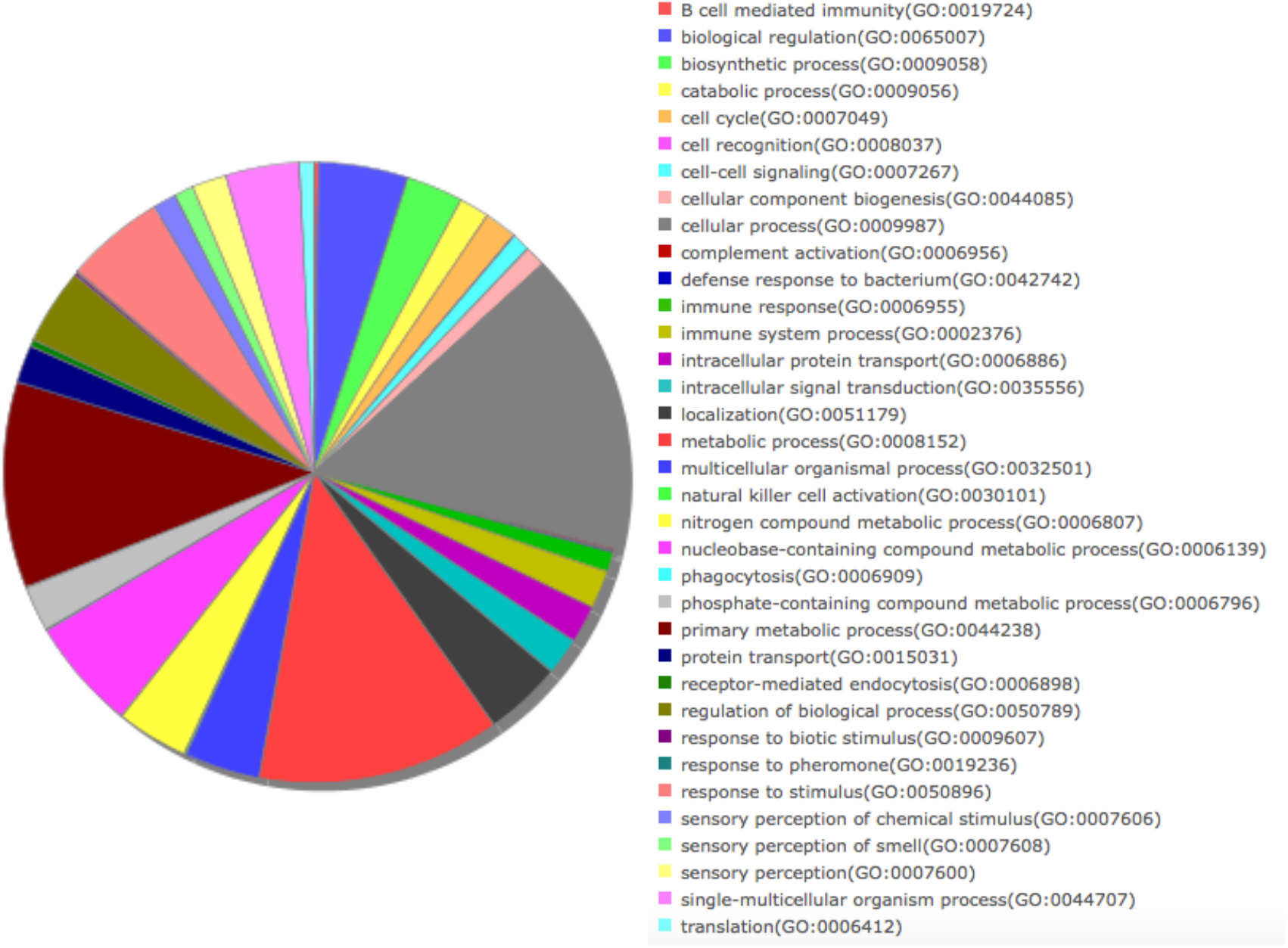
Functional annotation of the capybara genome with PANTHER.

**Fig S3.**
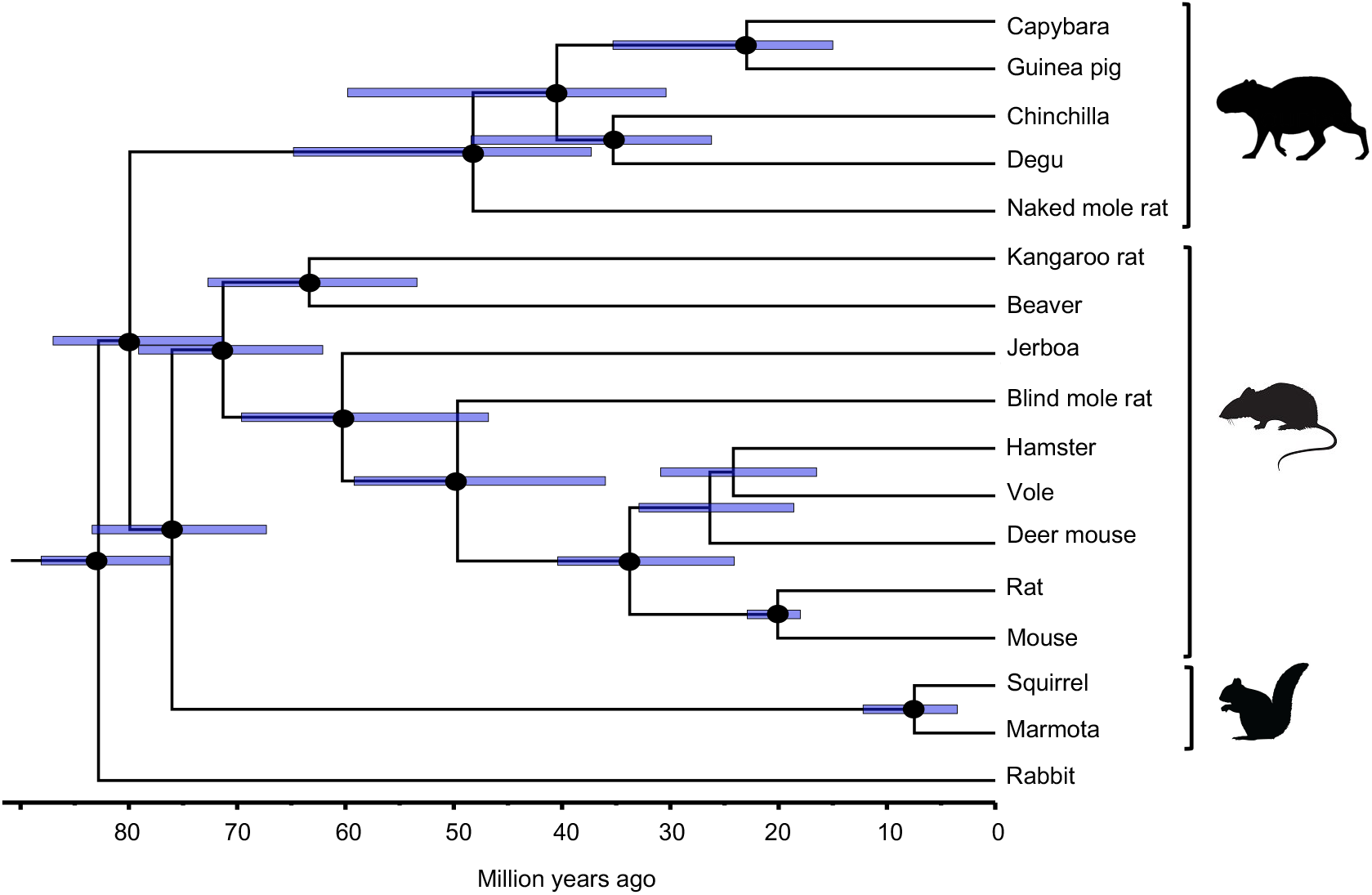
Divergence times of representative rodent species using the topology obtained from phylogenomic analyses. The blue bars on nodes represent the 95% credibility intervals of divergence time estimates. Two nodes were calibrated using fossil data as follows: Most recent common ancestor (MRCA) of Rodentia: 76 to 88 million years ago (Ma); MRCA of *Mus* and *Rattus:* 18 to 23 Ma. Black dots on nodes represent bootstrap values over 98% in both analyses: concatenated and species-tree. Clades from top to bottom: Hystricognathi, Mouse-related clade and Squirrel-related clade.

**Fig S4.**
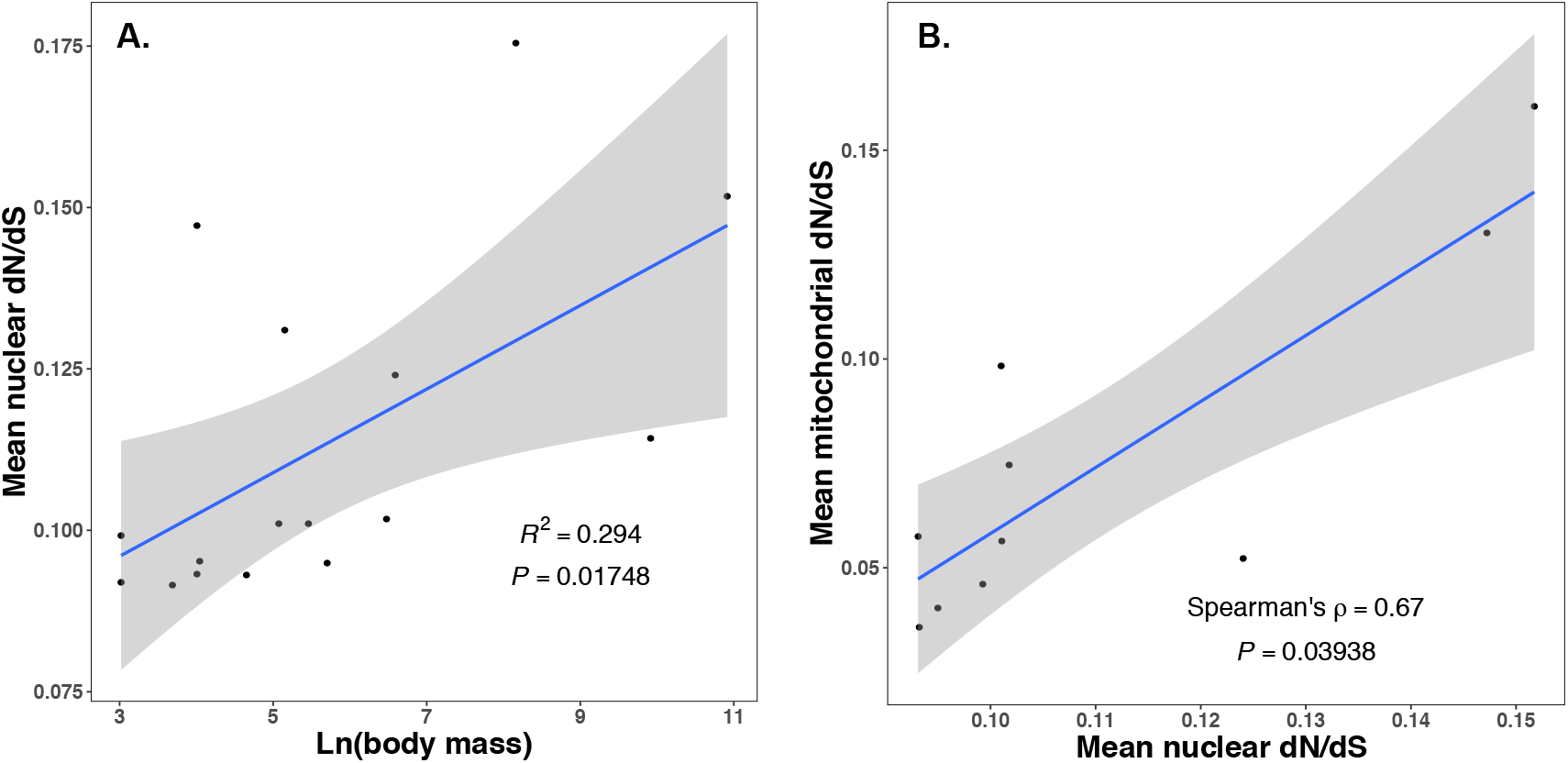
Demographic analyses of mutation load. **(A)** Correlation of log-transformed body mass values with the mean d_N_/d_S_ of 229 SCOs for each rodent species. Within rodents, mean d_N_/d_S_ is positively correlated with body mass indicating a lower efficiency of purifying selection relative to genetic drift in bigger rodents (R^2^ = 0.294, *df* = 14, P < 0.05; PIC: R^2^ = 0.332, *df* = 13, P < 0.05). **(B)** Correlation between the mean d_N_/d_S_ values of 229 SCOs and the mean dN/dS values of the 13-protein-coding mitochondrial genes for ten rodent species for which both nuclear and mitochondrial genomes were available. Nuclear and mitochondrial d_N_/d_S_ values show a significant positive correlation (Spearman’s ρ = 0.67, P < 0.05; PIC: Spearman’s ρ = 0.7, P < 0.05), which indicates a reduction of the effective population size caused by the increment in body mass.

**Fig S5.**
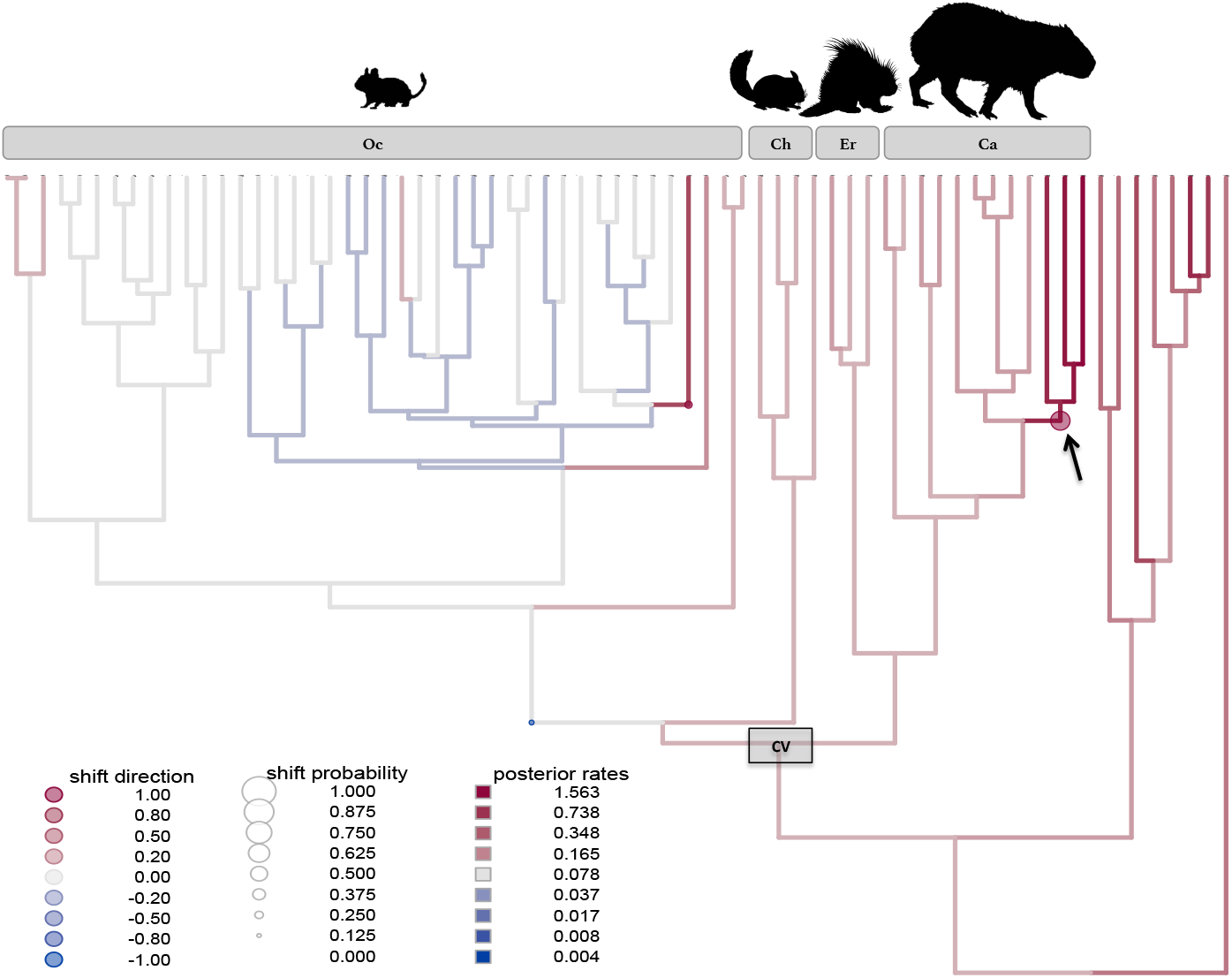
Bayesian sampling of evolutionary rates of body mass evolution. Shown is a significant upturn in the evolutionary rate of this trait in the most recent common ancestor of the clade containing capybara-rock cavy-mara (arrow). Hue and size of circles denote posterior support of a rate shift in the branch, with larger and redder circles suggesting higher posterior probability supporting an upturn in the evolutionary rate. Branch color denotes posterior estimated rates of body mass evolution. CV: Caviomorpha, Oc: Octodontoidea (degus and spiny rats), Ch: Chinchilloidea (chinchillas and viscachas), Er: Erethizontoidea (porcupines), Ca: Cavioidea (capybaras, maras and guinea pigs).

**Fig S6.**
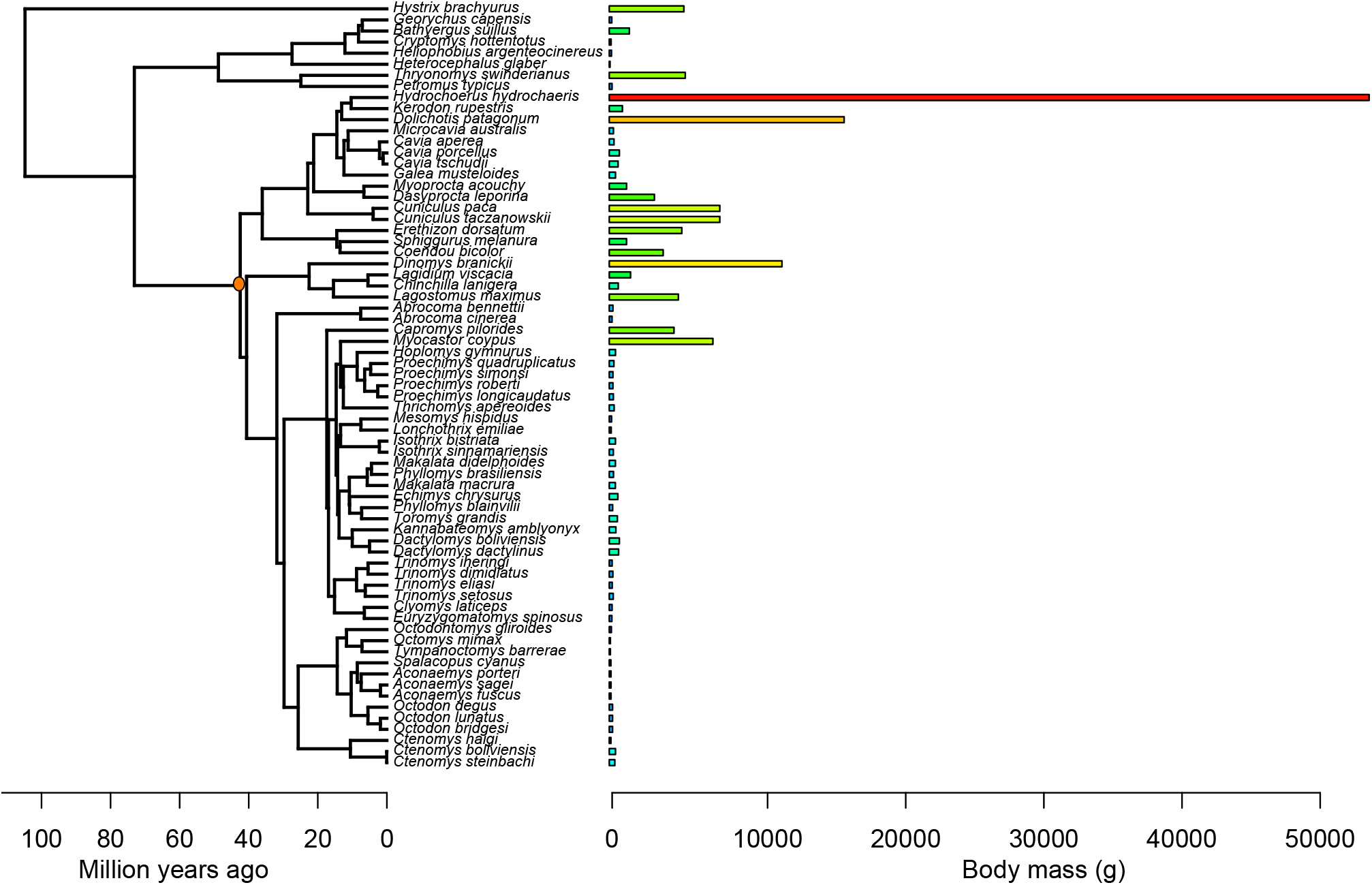
Time-calibrated phylogram of caviomorph rodents (from [100]) with values of body mass indicated to the right of each species. The broad range in body mass within caviomorph rodents reaches three orders of magnitude, with the smallest species weighting under 100g (green bars) and the largest species, the capybara, weighting 55,000g (red bar). The red node corresponds to the most recent common ancestor of caviomorph rodents.

